# Overexpression of TopBP1 leads to transformation with a TP53 mutation of non-tumorigenic breast epithelial cells

**DOI:** 10.1101/2022.04.05.487132

**Authors:** Rintu M Umesh, Mayurika Lahiri

## Abstract

DNA topoisomerase IIβ - binding protein 1 (TopBP1) is a mediator protein that regulates the cell cycle checkpoint signaling pathway. A plethora of studies suggests high TopBP1 levels are positively associated with various cancers. Although TopBP1 transcript, as well as protein expression levels, are high in breast cancers, its role in breast tumorigenesis is not yet explored. In our studies, we observed that TopBP1 levels are high in premalignant and malignant cells of the MCF10A cancer progression series compared to the non-tumorigenic MCF10A cells. In order to establish the role of TopBP1 in tumorigenesis, TopBP1 overexpression in non-tumorigenic MCF10A, and stable knock-down in malignant MCF10CA1a cells were performed and grown in Matrigel^™^ as breast spheroids.

Overexpression of TopBP1 in MCF10A spheroids induced hyperproliferation, disruption of polarity and cell-cell junctions. Moreover, TopBP1 overexpressing 3D dissociated cells exhibited EMT-like phenotype and tumorigenic properties such as increased cell migration, invasion, colony formation capabilitiy and anchorage-independent growth, indicating acquisition of cellular transformation. Finally, we demonstrated TopBP1 overexpressing cells to form tumors in athymic mice thereby confirming their tumorigenic potential. We also confirmed that overexpression of TopBP1 led to a mutation in TP53 and other genomic insults. To summarise, we observed that ectopic expression of TopBP1 transforms MCF10A breast epithelial cells. These transformed cells harbour phenotypic and genotypic characteristics similar to that of malignant cells.

## Introduction

DNA topoisomerase IIβ-binding protein 1 (TopBP1) participates in the activation of the DNA damage checkpoint to maintain genome integrity (Xu and Leffak, 2010). TopBP1 gene is located at chromosome 3q22.1, and is conserved across species (Wardlaw et al., 2014). It was first identified through two independent screens in *S. pombe* as rad4 (Schupbach, 1971), and cut5 (Hirano et al., 1986). Further human TopBP1 was identified as an interactor of Topoisomerase IIβ in a yeast two-hybrid screen (Wang and Elledge, 2002). The other known homologues of TopBP1 are Dpb11 (*S. cerevisiae*) (Araki et al., 1995), Mus101 (*D. melanogaster*) (Yamamoto et al., 2000), Mei1 (*A. thaliana*) and Xmus101 (*X. laevis*).

TopBP1 has eight BRCA1 C-terminus (BRCT) repeat domains through which it interacts with other proteins (Yu et al., 2003) and an ATR activation domain (AAD) which is necessary to stimulate the kinase activity of ATR (Kumagai et al., 2006) and thus checkpoint activation. Other than its essential role in DNA damage response, TopBP1 is also involved in transcriptional regulation of genes involved in cell proliferation and apoptosis such as E2F1 and c-Myc (Liu et al., 2003). Akt phosphorylates TopBP1 at serine residue 1159 which leads to its oligomerization through the BRCT domains 7 and 8 and thus shifts its function from checkpoint activation to transcriptional regulation (Liu et al., 2006). Oligomerized TopBP1 binds to E2F1 and inhibits E2F1-mediated apoptosis (Liu et al., 2006). TopBP1’s interaction with BLM (Bloom syndrome helicase) has been reported to inhibit sister chromatid exchange thus maintaining genome stability (Wang et al., 2013). TopBP1 also plays a role during replication initiation by interacting with Treslin to facilitate Cdc45 loading onto the replication origins (Kumagai et al., 2006). At the physiological level, TopBP1 is required for the G1 to S transition in the cell cycle.

Aberrant expression of TopBP1 has been observed in breast cancers. Immunohistochemical studies performed on breast cancer biopsy samples showed TopBP1 to be upregulated in breast cancer tissues. TopBP1 overexpression was associated with higher-grade tumors and reduced patient survival (Forma et al., 2012; Going et al., 2007; Liu et al., 2009). p53 is a well-studied tumor suppressor and in the majority of cancers, p53 is mutated leading to gain of function mutations in the protein.TopBP1 interacts with the DNA binding domain of p53 through its BRCT domains 7 and 8 and thus inhibits the promotor binding activity of p53 (Liu et al., 2009). In their study, Liu and colleagues showed that TopBP1 overexpression reduced the p53 transcriptional activity upon DNA damage. However, the levels of total p53 and its activation at the serine residue 15 remained unaffected (Liu et al., 2009). TopBP1 interacts with p53 hot spot mutants and increase proliferation and decrease apoptosis by affecting p53 transcriptional activity (Liu et al., 2011). Other studies have shown TopBP1 to induce chemoresistance by upregulating p53 levels (Lv et al., 2016). PI(3)K/Akt, p53 and E2F1 are some of the common pathways that are deregulated in cancers (Cancer Genome Atlas, 2012). A recent study found that TopBP1 paradoxically hinders ATR/Chk1 activation in cancers (Liu et al., 2021). As mentioned previously, TopBP1 regulates the function of various cancer regulatory proteins such as PI(3)K/Akt, p53 and E2F1 and affects the viability of cancer cells making it a novel therapeutic target (Chowdhury et al., 2014).

There are a number of studies that suggest an oncogenic role for TopBP1. Depletion of TopBP1 makes various cancer cells susceptible to undergo apoptosis, however, higher levels of TopBP1 promotes the survival of cancer cells. Most of the published studies focus and emphasize on the tumor-promoting property of TopBP1. To investigate the role of TopBP1 in breast tumorigenesis, the MCF10A cancer progression series was used as a model system. The MCF10A series consists of MCF10A (non-transformed epithelial cells), MCF10AT1 (premalignant cells) and MCF10CA1a (malignant cells). MCF10A was derived from the parental MCF10 cells from a woman with fibrocystic mammary tissue (Soule et al., 1990). The adherent MCF10A cells were transformed with constitutive H-RAS to generate MCF10AT1. MCF10CA1a was derived by continuous passaging of MCF10AT1 in mice for 100 generations (Santner et al., 2001). In our study, we report a novel tumor initiatory role of TopBP1. In this study, we demonstrate that ectopic expression of TopBP1 in non-tumorigenic cells promote the signaling cascade in favor of initiation of various tumorigenic properties. Breast epithelial cells with ectopic expression of TopBP1 when grown as 3D cultures formed spheroids with increased size, filled lumen, hyperproliferation, and altered polarity similar to spheroids formed by malignant cells. Also, the cells dissociated from the 3D cultures of TopBP1 overexpressing acini had morphology similar to the malignant MCF10CA1a cells. As expected, knockdown of TopBP1 in the malignant cells represses the spheroidal size and proliferation in the 3D cultures. TopBP1 overexpressing cells dissociated from 3D cultures exhibited increased migration, invasion, colony formation, anchorage-independent growth and tumorigenic potential similar to the malignant MCF10CA1a cells. TopBP1 overexpression in non-tumorigenic cells induced endogenous DNA damage and activation of DNA damage response (DDR) similar to the malignant cells suggesting its role in maintaining genome stability. TopBP1 overexpressing cells also exhibited mutation in TP53 gene similar to the malignant MCF10CA1a cells. Taken together we report TopBP1 as a novel initiator as well as a regulator of breast tumorigenesis that is required to maintain genomic stability.

## Results

### Overexpression of TopBP1 leads to an increase in breast acinar size

To investigate the role of TopBP1 in the transformation of breast epithelial cells, TopBP1 protein expression levels were analyzed in the MCF10A cancer progression series. There was a significant increase in TopBP1 expression in pre-malignant MCF10AT1 with a further increase in the malignant MCF10CA1a cells when compared to the non-tumorigenic MCF10A cells (Figure 1A-C). These results suggest TopBP1 may play an important role during the process of tumorigenesis. To investigate further the role of TopBP1 in breast tumorigenesis, TopBP1 was overexpressed in MCF10A (TopBP1 OE) (Figure 1D-E), and the transformative potential of these cells was analyzed. TopBP1 OE cells formed large spheroids similar to multiacinar cultures with filled lumen as compared to the control cells (mCherry-MCF10A) (Figure 1F). There was a significant increase in the surface area and volume of the TopBP1 OE spheroids when compared to the control acini (Figure 1G-H). Interestingly, TopBP1 OE spheroids showed a reduction in sphericity when compared to the control acini suggesting the presence of protrusion-like phenotype that are the initial events of transformation (Figure 1I). TopBP1 OE spheroids were dissociated and grown as monolayer cultures. TopBP1 OE 3D dissociated cells exhibited morphology similar to the malignant MCF10CA1a cells (Figure 1J). However, when TopBP1 OE cells were grown as monolayer cultures, they showed phenotypes similar to the non-tumorigenic MCF10A cells (Figure 1K). These results suggest that overexpression of TopBP1 in non-tumorigenic MCF10A cells grown as three-dimensional breast acinar cultures leads to morphological changes that are similar to the malignant cells MCF10CA1a.

**Figure 1:**
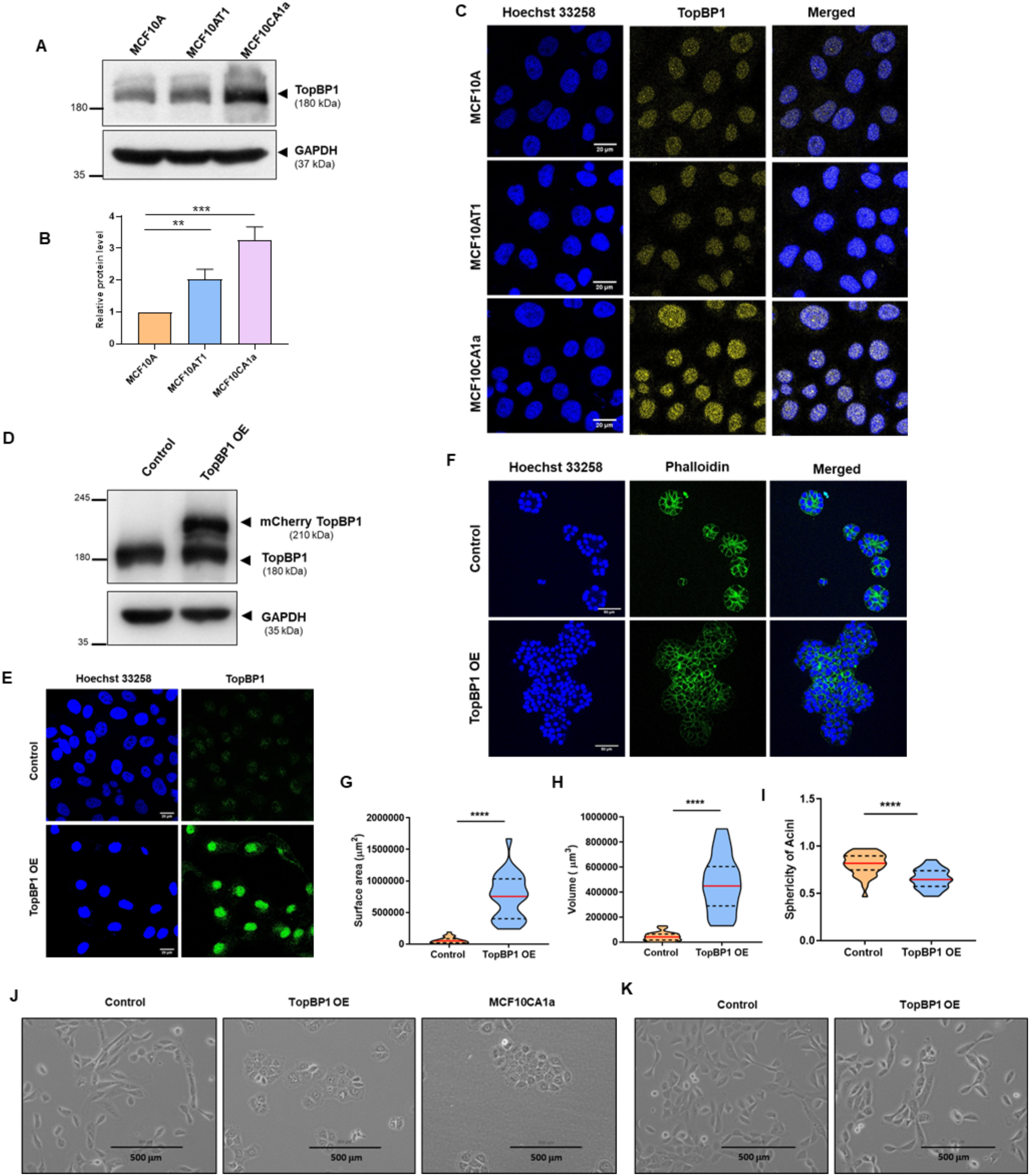
TopBP1 overexpression leads to increased acinar size in MCF10A 3D cultures. A) Western blot analysis and C) immunofluorescence assay showing varying levels of TopBP1 in MCF10A, MCF10AT1 and MCF10CA1a (2D) cells. GAPDH was used as loading control. B) Quantification showing the fold change of TopBP1 levels after normalizing with GAPDH. Statistical analysis was done using One-way ANOVA (*** p <0.001, **p <0.01). D) MCF10A cells stably expressing mCherry (control) and TopBP1 OE (2D) were lyzed and immunoblotting was performed using TopBP1 specific antibody. E) mCherry MCF10A (control) and TopBP1 OE MCF10A (2D) cells were immunostained using TopBP1 specific antibody to confirm the overexpression (scale bar: 20 μm). F) Representative images of mCherry MCF10A (control) and TopBP1 OE MCF10A cells grown as 3D “on-top” cultures for 10 days and fixed and stained for phalloidin (green) and Hoechst 33258 (blue) (scale bar: 50 μm). G) Surface area, H) Volume, and I) Sphericity of TopBP1 OE MCF10A acini were calculated using Huygens software (SVI, Netherlands) and represented as violin plots. >50 acini from 3 independent experiments were analyzed. Statistical analysis was done using the Mann Whitney test (**** p <0.0001). J) Representative images of mCherry MCF10A (control) and TopBP1 OE MCF10A 3D dissociated and MCF10CA1a 2D cells. K) Representative images of mCherry and TopBP1 OE MCF10A 2D cells. Images were acquired on the ECLIPSE TS100 Nikon microscope (scale bar: 500 μm).

### TopBP1 knockdown reduces spheroid size in malignant cells

As we have demonstrated MCF10CA1a to have higher levels of TopBP1 and TopBP1 OE cells morphologically resembled MCF10CA1a cells, we next investigated whether stable knockdown of TopBP1 in MCF10CA1a would lead to a reversal from the malignant to the non-tumorigenic phenotype when grown as 3D cultures (Figure 2A-C). Filled lumen and lower sphericity are properties associated with cancer cells. Upon TopBP1 knockdown, MCF10CA1a cells formed spheroids with a clear lumen (Figure 2D). The surface area and volume of the spheroids of shTopBP1-MCF10CA1a spheroids were significantly reduced when compared to the control pLKO.1 MCF10CA1a spheroids (Figure 2E-F). The sphericity of these spheroids increased upon TopBP1 KD (Figure 2G). These results suggest that TopBP1 KD in MCF10CA1a reduces spheroid size, thus leading to alteration in the morphology of the malignant cells.

**Figure 2:**
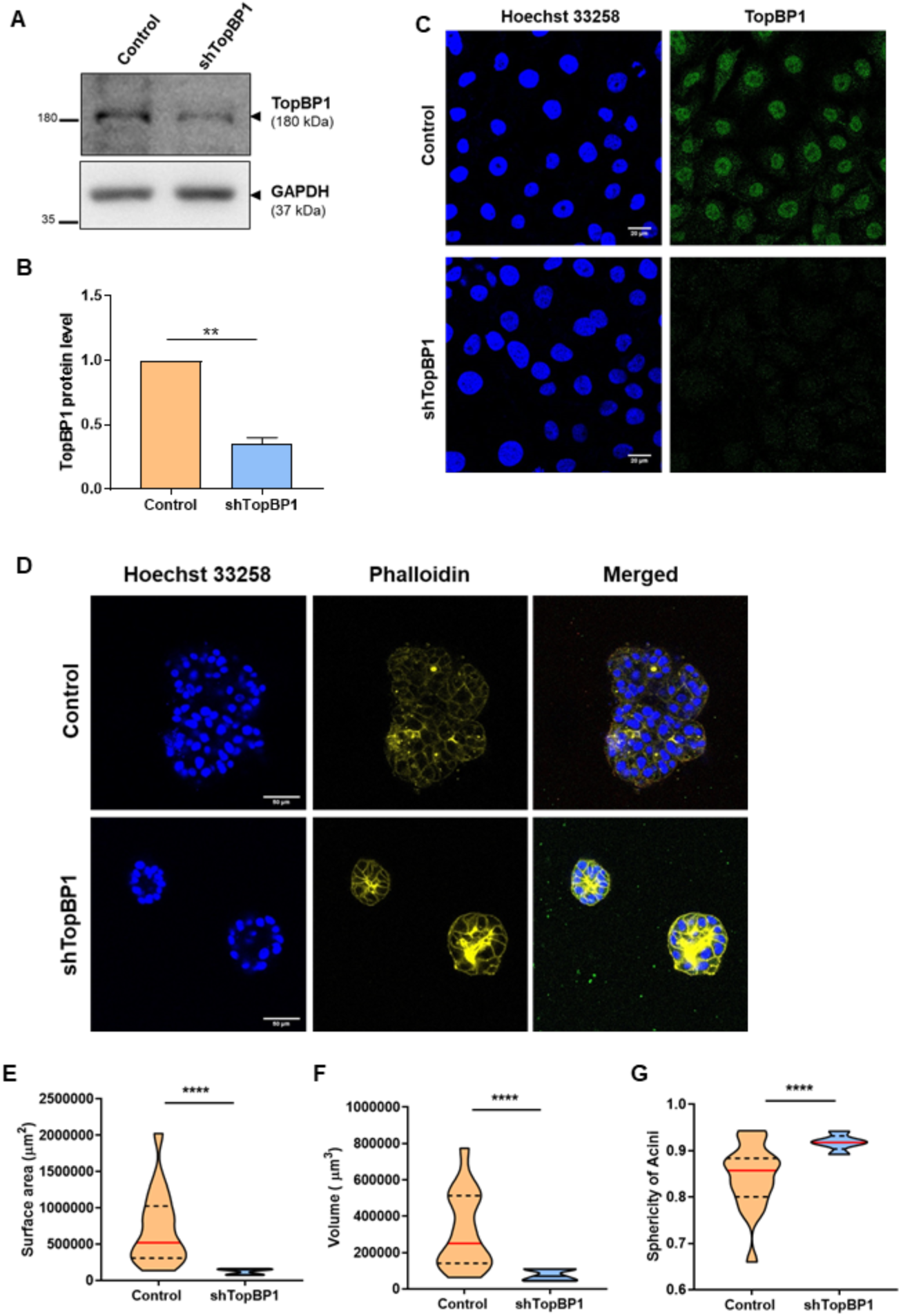
TopBP1 knockdown reduces spheroid size in MCF10CA1a 3D cultures. MCF10CA1a cells stably expressing pLKO1.TRC (control) and shTopBP1 (2D) were A) lyzed and immunoblotting was performed using TopBP1 specific antibody. B) The fold change of TopBP1 was quantified after normalizing with GAPDH. Statistical analysis was done using Student’s t-test (** p <0.01). C) pLKO.1 MCF10CA1a (control) and shTopBP1 MCF10CA1a 2D cells were immunostained using TopBP1 specific antibody to confirm the knockdown (scale bar: 20 μm). D) Representative images of pLKO.1 MCF10CA1a (control) and shTopBP1 MCF10CA1a cells grown as 3D cultures for 10 days and fixed and stained for phalloidin (yellow) and Hoechst 33258 (blue) (scale bar: 50 μm). E) Surface area, F) Volume and G) Sphericity of these acini were calculated using Huygens software (SVI, Netherlands) and represented as violin plots. >50 acini from 3 independent experiments were analyzed. Statistical analysis was done using Mann Whitney test (**** p <0.0001).

### Overexpression of TopBP1 leads to cell hyperproliferation

Our results demonstrate that acinar size increases upon overexpression of TopBP1. To further corroborate this data, the number of cells that form the acini were calculated. MCF10A acini are normally comprised of 30-35 cells. Overexpression of TopBP1 led to a significant increase in the percentage of acini that were comprised of more than 50 cells when compared to the control (Figure 3A). Upon knockdown of TopBP1 in MCF10CA1a, there was a significant reduction in the percentage of acini that had more than 50 cells when compared to the control (Figure 3B). These results suggest TopBP1 expression levels affect proliferation. To investigate the proliferative potential of TopBP1, acini ectopically expressing TopBP1 were stained for Ki67, a proliferation marker. It was observed that 100% of TopBP1 OE acini had more than 33% Ki67 positive cells when compared to the control (Figure 3C) which was similar to the malignant MCF10CA1a spheroids (Figure 3C). Immunoblotting analysis demonstrated increased Ki67 protein expression in the lysates from TopBP1 OE and malignant MCF10CA1a spheroids when compared to the control (Figure 3D-E). TopBP1 knockdown led to a significant reduction in Ki67 expression levels in MCF10CA1a 3D lysates (Figure F-G). These results suggest TopBP1 to regulate proliferation of cells grown as 3D cultures. TopBP1 OE 3D dissociated cells also exhibited increased proliferation when compared to the control as was observed by an increase in cell numbers (Figure 3H). This increase in proliferation indicates a higher fraction of the cells in the S and G2-M phases of the cell cycle. To investigate the number of cells in each phase of the cell cycle, the cell cycle profile of control, TopBP1 OE and MCF10CA1a cells were analyzed. It was observed that TopBP1 OE had a higher percentage of cells in the S (14%) and G2-M (31%) phases, which was similar to the malignant MCF10CA1a cells (S: 13% and G2-M: 42%) when compared to the control (S: 7% and G2-M: 16%) (Figure 3I). Cells were also analyzed for the expression of two other S phase cell cycle markers, PCNA and Cyclin A. Both PCNA and Cyclin A expression levels were significantly higher in TopBP1 OE cells when compared to the control. This increase in expression was similar to the expression pattern of both PCNA and Cyclin A in MCF10CA1a cells (Figure 3J-O). These results conclude that overexpression of TopBP1 leads to cellular hyper-proliferation, which is one of the hallmarks of cancer cells.

**Figure 3:**
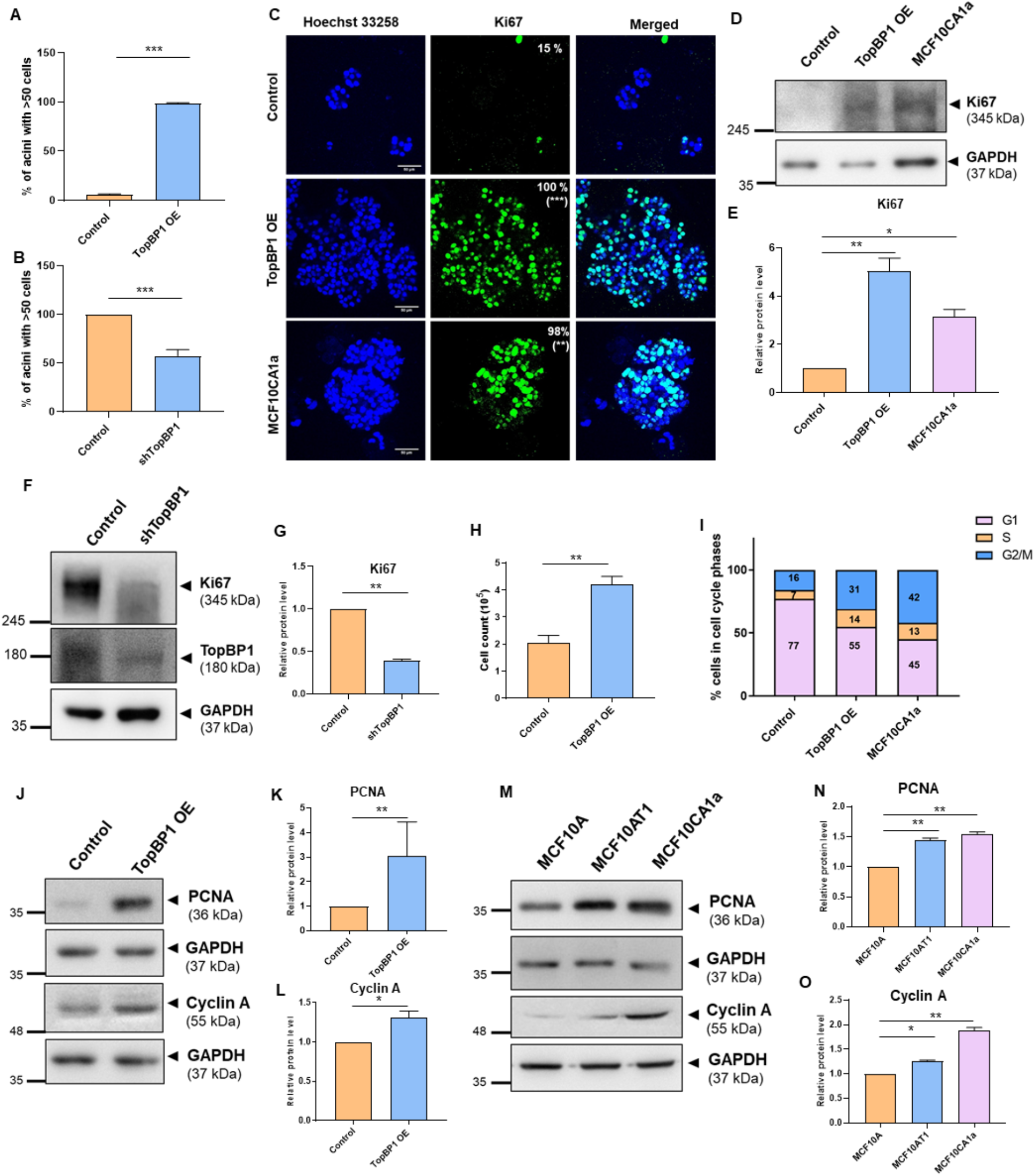
TopBP1 overexpression increases proliferation. The number of cells per acini of A) mCherry MCF10A (control), TopBP1 OE MCF10A, B) pLKO1-MCF10CA1a (control) and shTopBP1 MCF10CA1a were counted manually and the percentage of acini with more than 50 cells were plotted as a bar graph. C) Representative images of mCherry MCF10A (control), TopBP1 OE and MCF10CA1a cells grown as 3D cultures for 10 days and fixed and immunostained for Ki67 (green) (scale bar: 50 μm). Number of Ki67 positive cells were manually counted and divided by total number of cells per acini. Percentage of acini with more than 33% of Ki67 positive cells (clinically relevent value) were mentioned on the upper right corner of the Ki67 image. >50 acini from 3 independent experiments were analyzed. Statistical analysis was done using Mann Whitney test (** p <0.01, *** p <0.001). D) 16-day 3D cultures of mCherry MCF10A (control), TopBP1 OE spheroids and 10-day cultures of MCF10CA1a spheroids were lyzed and immunoblotting was performed using Ki67 specific antibody. E) Quantification showing the fold change of Ki67 after normalizing with GAPDH. F) 10-day 3D cultures of pLKO1-MCF10CA1a (control) and shTopBP1 MCF10CA1a were lyzed and immunoblotting was performed using Ki67 specific antibody. G) Quantification showing the fold change of Ki67 after normalizing with GAPDH. H) mCherry MCF10A (control) and TopBP1 OE MCF10A 3D dissociated cell number was counted using Scepter™ 72 hrs post-seeding. Statistical analysis was done using a Paired Student’s t-test (** p <0.01). I) mCherry MCF10A (control), TopBP1 OE MCF10A 3D dissociated cells and MCF10CA1a 2D cells were fixed in ethanol and stained using propidium iodide. Percentage of cells in the different cell cycle phases were analyzed on a BD Accuri flow cytometer and plotted as box plots. J) mCherry MCF10A (control) and TopBP1 OE MCF10A 3D dissociated cells and M) MCF10A, MCF10AT1 and MCF10CA1a 2D cells were lyzed and immunoblotting was performed to analyze the levels of PCNA and Cyclin A. Quantification showing the fold change of J, K, M and N) PCNA and J, L, M and O) Cyclin A after normalizing to GAPDH. Statistical analysis was done using Paired Student’s t-test for TopBP1 OE and One-way ANOVA for MCF10A series ( ** p <0.01, * p <0.05).

### TopBP1 overexpression leads to disruption of polarity

De-regulation of polarity is one of the characteristics of epithelial cancers (Debnath and Brugge, 2005). To investigate the effect of TopBP1 on polarity, TopBP1 OE acini were stained using basal polarity markers a6-integrin, and Laminin V. Loss or mislocalization of a6-integrin (Figure 4A) and Laminin V (Figure 4B) at the basal region were observed in TopBP1 OE speroids, similar to that of the malignant spheroids (Figure 4A-B). Golgi is apically located in MCF10A 3D cultures and is thus considered as an apical polarity marker (Debnath et al., 2003). To investigate whether TopBP1 plays a role in maintaining apical polarity, TopBP1 OE acini were stained with GM130, a cis-Golgi marker. It was observed that TopBP1 OE acini had dispersed Golgi architecture compared to the compact apically located Golgi in the control acini (Figure 4C). Golgi dispersal was further confirmed by staining the 3D dissociated TopBP1 OE cells using GM130. Similar dispersed Golgi was also observed in the malignant cells (Figure 4D), which was quantified by measuring the Golgi area (Figure 4E). β-catenin, a cell-cell junction marker, showed dispersed or speckle-like staining in TopBP1 OE acini when compared to the control acini, which was similar to MCF10CA1a spheroids suggesting disruption of cell-cell contact (Figure 4F). These results suggest that overexpression of TopBP1 led to disruption of the apicobasal and lateral polarity of MCF10A acini which is comparable to the disruption observed in the malignant MCF10CA1a spheroids, thus indicating a transformative potential of TopBP1.

**Figure 4:**
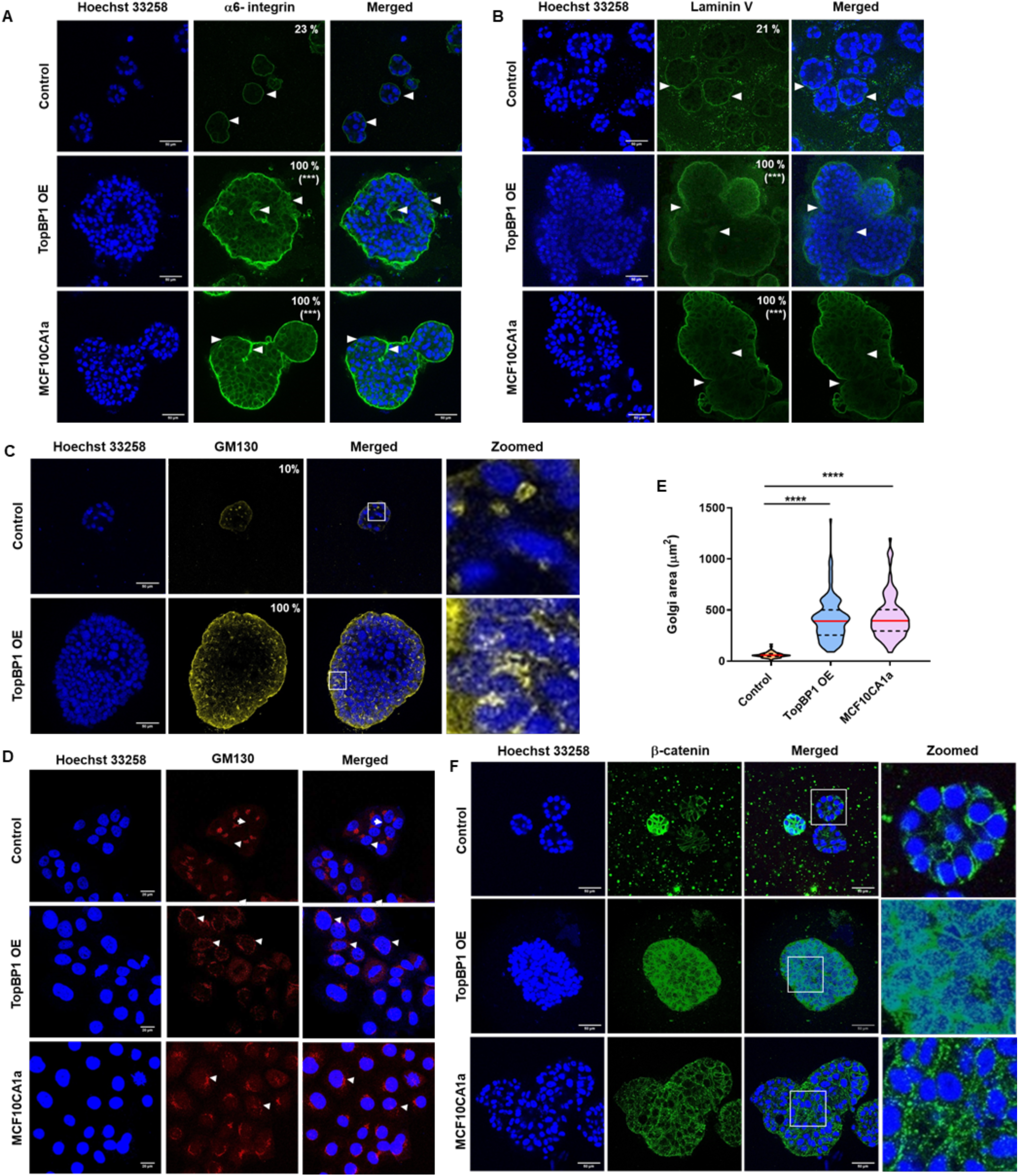
TopBP1 overexpression disrupts cell polarity. Representative images of mCherry MCF10A (control), TopBP1 OE MCF10A and MCF10CA1a cells grown as 3D cultures for 10 days, fixed and immunostained for A) a6-integrin B) Laminin V, C) GM130 and F) β-catenin (scale bar: 50 μm). D) mCherry MCF10A (control), TopBP1 OE MCF10A 3D dissociated and MCF10CA1a 2D cells were immunostained using GM130, a Golgi marker and E) Golgi dispersal was calculated by measuring Golgi area using ImageJ software. >50 acini from 3 independent experiments were analyzed. Statistical analysis was done using Mann Whitney test (*** p <0.001).

### Expression of EMT markers in TopBP1 overexpression is similar to malignant cells

Epithelial to mesenchymal transition (EMT) is a process acquired by cancer cells that favors their motility and invasiveness. To investigate the function of TopBP1 in the regulation of EMT, the expression of various EMT markers were analyzed. It was observed that there were acini with varying expressions of vimentin, a mesenchymal marker. 100% of mCherry-MCF10A acini (control) were vimentin-positive compared to only 11% of TopBP1 OE acini (Figure 5A). Similarly, only 14% of the MCF10CA1a 3D spheroids were vimentin-positive (Figure 5A). The TopBP1 OE dissociated cells also exhibited heterogeneity in vimentin expression similar to that of MCF10CA1a cells. Only 14% of TopBP1 OE and 23% of MCF10CA1a cells were vimentin-positive, whereas 100%of control cells were vimentin-positive (Figure 5B). Vimentin expression was further confirmed using immunoblotting where TopBP1 OE cells were observed to have negligible levels of vimentin when compared to the control, which was also similar to the low levels observed in MCF10CA1a cells (Figure 5C-D and S1A-B). The levels of the epithelial marker, E-cadherin and mesenchymal marker, N-cadherin, were also analyzed. TopBP1 OE cells had significantly higher levels of E-cadherin and reduced levels of N-cadherin when compared to the control (Figure 5E and S1C and E). The higher levels of E-cadherin and lower levels of N-cadherin observed in TopBP1 OE was similar to the observed levels in the malignant cells (Figure 5F and S1D and F). Other EMT markers like Slug, Twist, cytokeratin 14 and cytokeratin 19 were also analyzed in TopBP1 OE 3D dissociated cells and the pre-malignant and malignant cells. Slug and cytokeratin 14 levels were significantly reduced in TopBP1 OE, MCF10AT1 and MCF10CA1a lysates in comparison to the control, whereas Twist and cytokeratin 19 levels were observed to significantly upregulated (Figure 5G-H and S1G-N). These results demonstrate that overexpression of TopBP1 in the non-tumorigenic MCF10A cells steers the cells to acquire EMT-like characteristics similar to that of the malignant MCF10CA1a cells.

**Figure 5:**
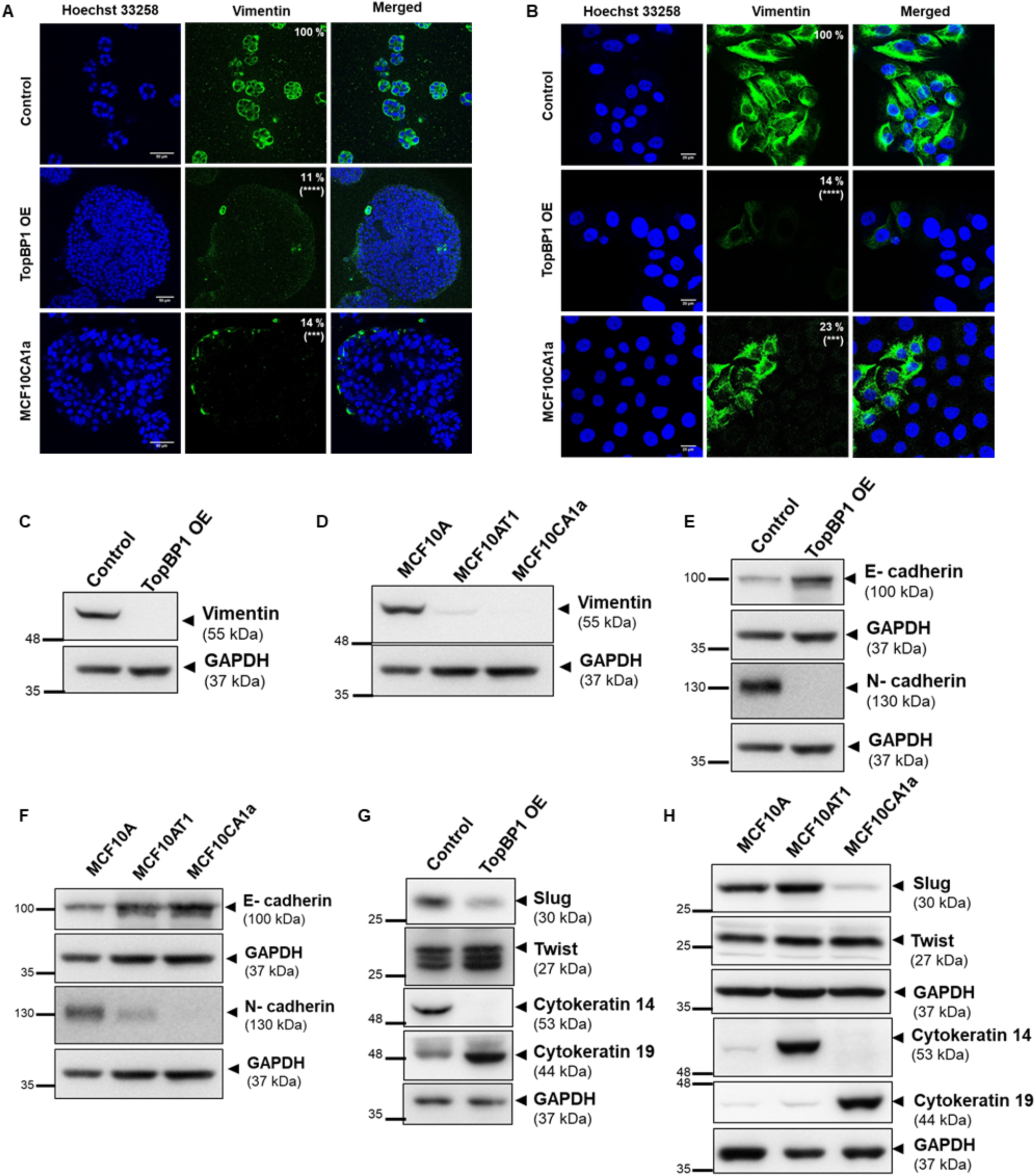
TopBP1 OE in MCF10A resembles MCF10CA1a EMT phenotypes. A) Representative images of mCherry MCF10A (control), TopBP1 OE MCF10A and MCF10CA1a cells grown as 3D cultures for 10 days, fixed and immunostained for vimentin. The number of vimentin-positive acini was manually counted and the percentage of positive acini was calculated and mentioned in the figures (scale bar: 50 μm). >50 acini from 3 independent experiments were analyzed. Statistical analysis was done using Mann Whitney test (*** p <0.001, **** p <0.0001). B) mCherry MCF10A (control) and TopBP1 OE MCF10A 3D dissociated and MCF10CA1a (2D) cells were immunostained using vimentin and the number of vimentin-positive cells were manually counted and the percentage of positive cells are mentioned in the figures. >300 cells from 3 independent experiments were analyzed. Statistical analysis was done using Mann Whitney test (*** p <0.001, **** p <0.0001). mCherry MCF10A (control) and TopBP1 OE MCF10A 3D dissociated cells as well as MCF10A, MCF10AT1 and MCF10CA1a (2D) cells were lyzed and immunoblotting was performed to analyze the levels of C-D) vimentin, E-F) E-cadherin, N-cadherin, G-H) Slug, Twist, Cytokeratin 14 and cytokeratin 19.

### TopBP1 overexpression leads to increased migration and invasion

In order for cancer cells to metastasize, the acquisition of both invasive and migratory potential is important. During invasion and migration, cells move either as single entities (Theveneau and Mayor, 2011) or collectively in small groups (Deisboeck and Couzin, 2009). Wound healing assays were performed to investigate collective cell migration of TopBP1 OE and MCF10CA1a cells. 100% wound closure was observed in TopBP1 OE and MCF10CA1a cells compared to 46% in the control cells after 12 hrs of creating the wounds thus confirming TopBP1 overexpression enabled the cells to undergo collective cell migration (Figure 6A-B). The single-cell migration potential of TopBP1 OE and MCF10CA1a cells were also investigated. TopBP1 OE cells showed increase in speed (Figure 6C), distance (Figure 6D) and displacement (Figure 6E) in comparison to the control cells. MCF10CA1a cells also showed increased speed and displacement compared to the control cells (Figure 6C-E). During the invasion process, invading cells secrete endopeptidases like matrix metalloproteinases (MMPs) into the surroundings to degrade the extracellular matrix proteins (Woessner, 1991). MMP2 and 9 are two such proteinases that have gelatinase activity. Gelatin zymography assay was performed to investigate whether TopBP1 overexpression leads to increased MMP secretion. It was observed that TopBP1 OE and MCF10CA1a 3D conditioned media had increased levels of MMP2 and MMP9 when compared to the control 3D media (Figure 6F-H). These results conclude that overexpression of TopBP1 in MCF10A cells leads to increased migration and invasion that is comparable to that observed in the malignant cells.

**Figure 6:**
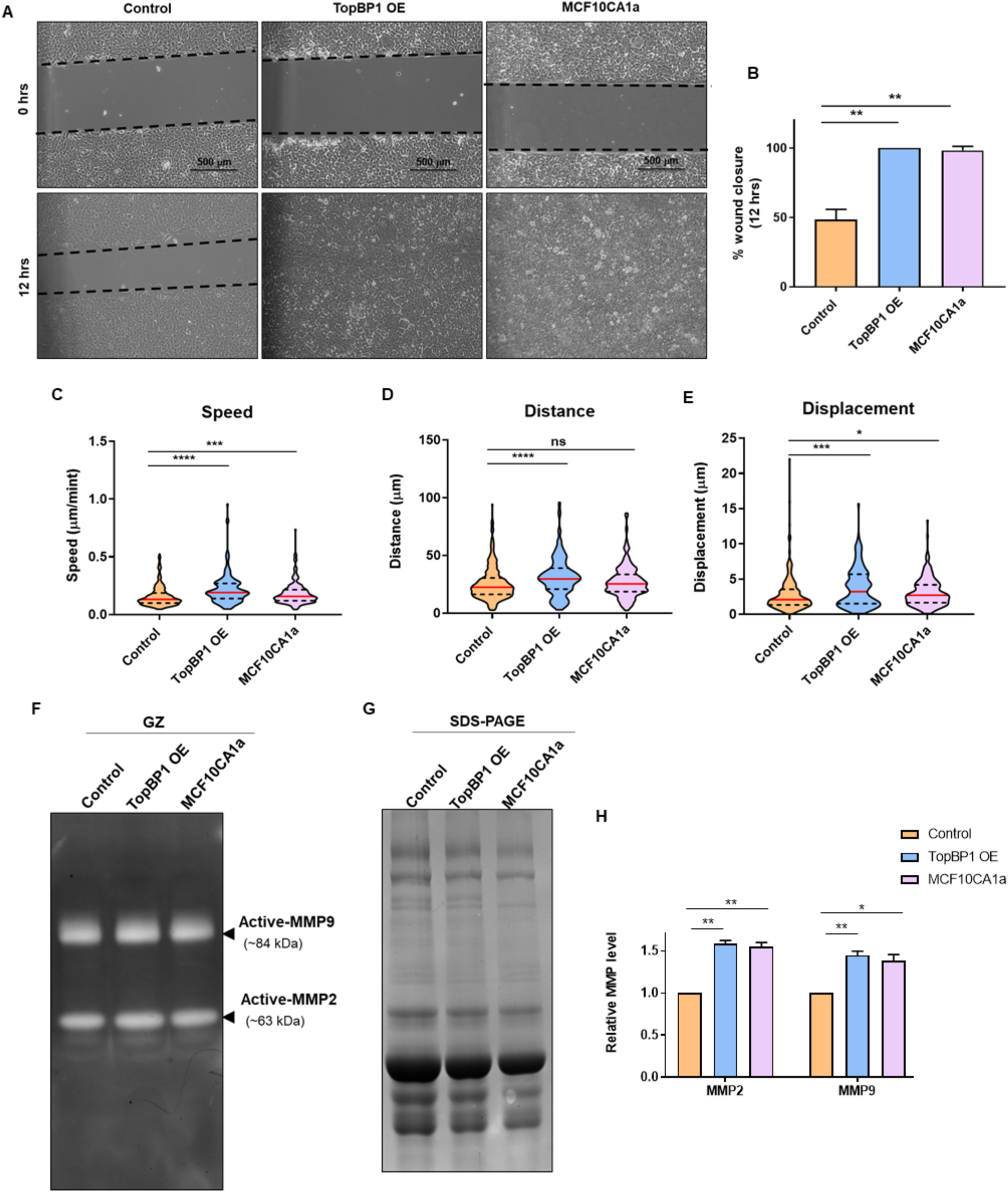
TopBP1 OE induces migration and invasion in MCF10A 3D dissociated cells. A) Representative images of wound healing assay demonstrating collective cell migration of mCherry MCF10A (control), TopBP1 OE MCF10A 3D dissociated cells and MCF10CA1a (2D) cells at 0 hrs and 12 hrs. Images were acquired on an ECLIPSE TS100 Nikon microscope at 10X magnification (scale bar: 500 μm). B) Wound area was manually measured using ImageJ and the percentage wound closure at 12 hrs was calculated and represented as a bar graph. Time-lapse imaging of mCherry MCF10A (control), TopBP1 OE MCF10A 3D dissociated cells and MCF10CA1a (2D) cells seeded as sparse populations were performed for a period of 3 hrs with images taken at 2-minute intervals on an Evos FL Auto microscope (Life Technologies). Cells were tracked using Fast track software and the C) speed, D) distance and, E) displacement were calculated and represented as violin plots. Gelatin zymography was performed using conditioned media from 10-day 3D cultures of mCherry MCF10A (control), TopBP1 OE MCF10A and MCF10CA1a. F) Gelatin zymography gel showing clearance of gelatin at the size of active MMP9 (~84 kDa) and MMP2 (~67 kDa) and G) the total protein in the media was quantified using SDS-PAGE. H) The MMP levels were quantified after normalizing to the total protein content of each sample. All data represented as Mean ± SEM from 3 independent biological experiments. Statistical analysis was done using One-way ANOVA (**** p <0.001, *** p <0.001, ** p <0.01, * p <0.05, ns p>0.05).

### TopBP1 overexpression induces transformation of MCF10A cells

In order to establish the role of TopBP1 in transformation, the colony formation ability of TopBP1 OE cells was studied. It was observed that TopBP1 OE cells formed colonies similar to that of the malignant cells while control cells failed to form colonies after 7 days (Figure 7A-B). Anchorage-independent growth is one of the properties of transformed cells. TopBP1 OE 3D dissociated cells were layered on soft agar to investigate their ability to grow in an anchorage-independent manner. TopBP1 OE and MCF10CA1a cells formed large colonies in 7 days, whereas the control cells failed to form colonies on soft agar (Figure 7C-D). To further establish the tumorigenic potential of TopBP1, TopBP1 OE and control cells were injected subcutaneously into the left and right flanks of athymic nude mice, respectively. Cells overexpressing TopBP1 formed tumors in the right flank within 4 weeks suggesting the tumor initiating role of TopBP1, whereas the control cells failed to form tumors even after 7 weeks. (Figure 7E-F). These results conclude that TopBP1 overexpression induces transformation of non-tumorigenic MCF10A cells that is capable of inducing tumorigenesis in mice.

**Figure 7:**
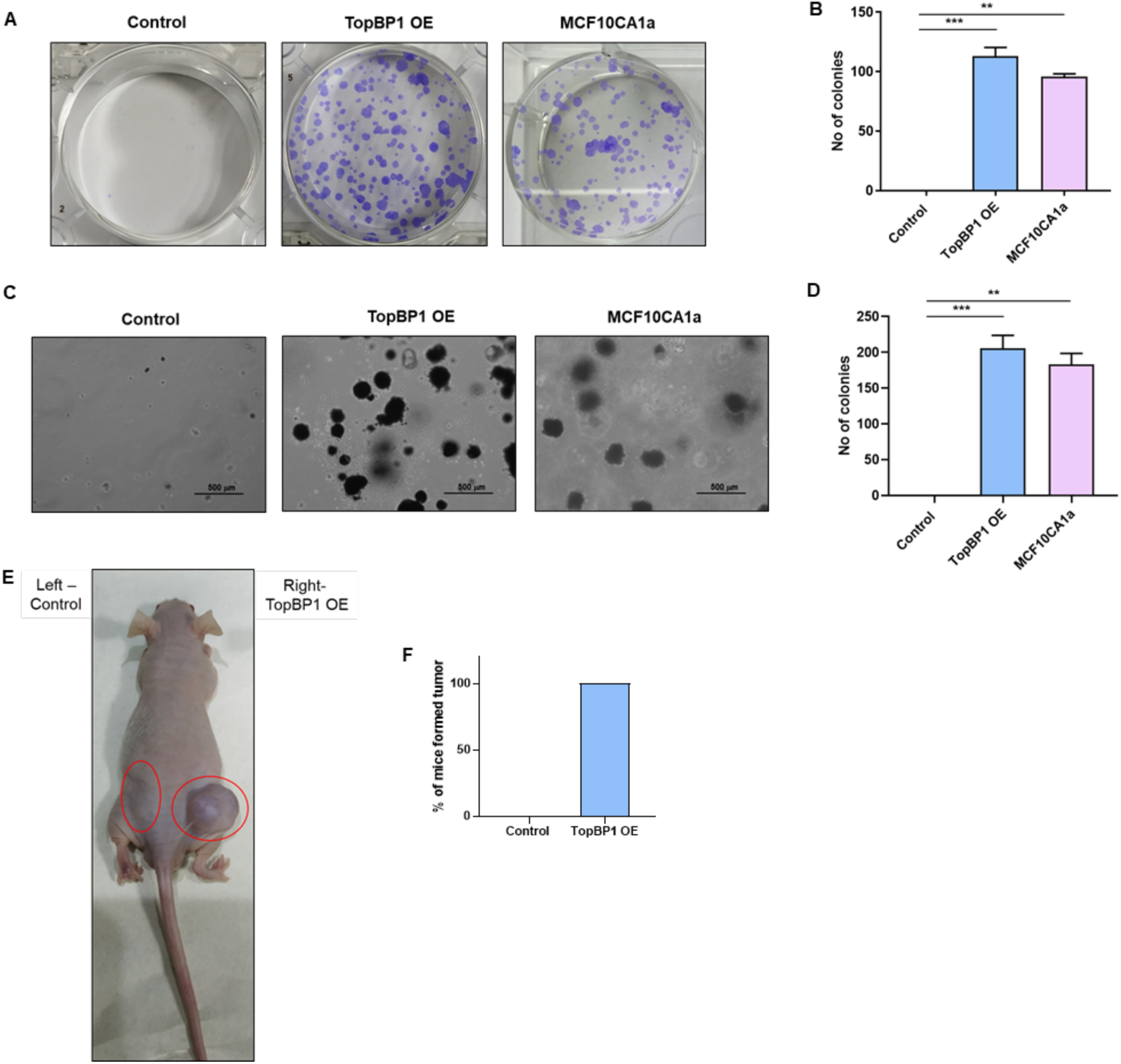
MCF10A cells acquire transformation potential upon TopBP1 OE in 3D cultures. A) Crystal violet stained colonies of mCherry MCF10a (control), TopBP1 OE MCF10A 3D dissociated and MCF10CA1a (2D) cells grown on cell culture-treated 35 mm culture plates for 7 days. B) The number of colonies were counted manually and represented as bar graphs. C) MTT-stained colonies of mCherry MCF10A (control), TopBP1 OE MCF10A 3D dissociated and MCF10CA1a (2D) cells grown on agar-coated 35 mm culture plates for 7 days. Images were acquired on an ECLIPSE TS100 Nikon microscope at 10X magnification (scale bar: 500 μm). D) Number of colonies were counted manually and represented as bar graphs. All data represented as Mean ± SEM from 3 independent biological experiments. Statistical analysis was done using One-way ANOVA (**** p <0.001, *** p <0.001, ** p <0.01, * p <0.05, ns p>0.05). E) Mice injected with mCherry MCF10A (control, left) and TopBP1 OE (right) 3D dissociated cells subcutaneously in flanks were sacrificed after 7 weeks of injection. F) Percentage of mice that formed tumor were represented as bar graph. Data pooled from N=12 mice.

### Overexpression of TopBP1 induces DNA damage that leads to activation of the DDR response

Genomic instability is one of the hallmarks of cancer (Hanahan and Weinberg, 2011). Defects in checkpoint activation can lead to the accumulation of DNA damage, leading to genomic instability (Kastan and Bartek, 2004). TopBP1 is a mediator protein in the DNA damage response (DDR) pathway and plays a pivotal role in the DDR and repair pathways. Our data confirms that overexpression of TopBP1 leads to the transformation of breast epithelial cells into a malignant phenotype. Thus it was intriguing to check whether overexpression of TopBP1 leads to an alteration in the cell’s response to DNA damage. To investigate whether TopBP1 overexpression induces DNA damage, we analyzed the activation of RPA (marker for single-strand breaks) and γH2AX (marker for double-strand breaks) in TopBP1 OE 3D dissociated cells. Interestingly, activation of RPA and γH2AX were observed in TopBP1 OE cells even in the absence of any external DNA damage (Figure 8A and S2 A-B). A similar pattern of activated pRPA and γH2AX were observed in the malignant MCF10CA1a cells compared to MCF10A and MCF10AT1, suggesting the presence of endogenous DNA damage in these transformed cells (Figure 8B and S2 C-D). As expected, knockdown of TopBP1 in the malignant MCF10CA1a cells showed relatively low activation of RPA and γH2AX (Figure 8C and S2 E-F). These results indicate that TopBP1 help to regulate the endogenous DNA damage inside the cells. Overexpression of the protein leads to an imbalance where TopBP1 is no longer able to perform its required function. We next investigated whether the endogenous DNA damage induced due to overexpressionTopBP1 leads to activation of the DNA damage response. As TopBP1 is a mediator protein in the single-strand DNA break pathway we analyzed the activation of ATR and Chk1 in TopBP1 OE cells. Both ATR and Chk1 were observed to be activated in TopBP1 OE cells compared to control cells. (Figure 8E and S2 G-H). A similar activation pattern of ATR and Chk1 was also observed in MCF10CA1a cells in comparison to the non-tumorigenic MCF10A cells (Figure 8F and S2 I-J). The knockdown of TopBP1 in the malignant cells led to reduced ATR and Chk1 activation (Figure 8G and S2 K-L). These results suggest that overexpression of TopBP1 leads to increased endogenous DNA damage and altered DNA damage response in MCF10A cells along with cellular transformation.

**Figure 8:**
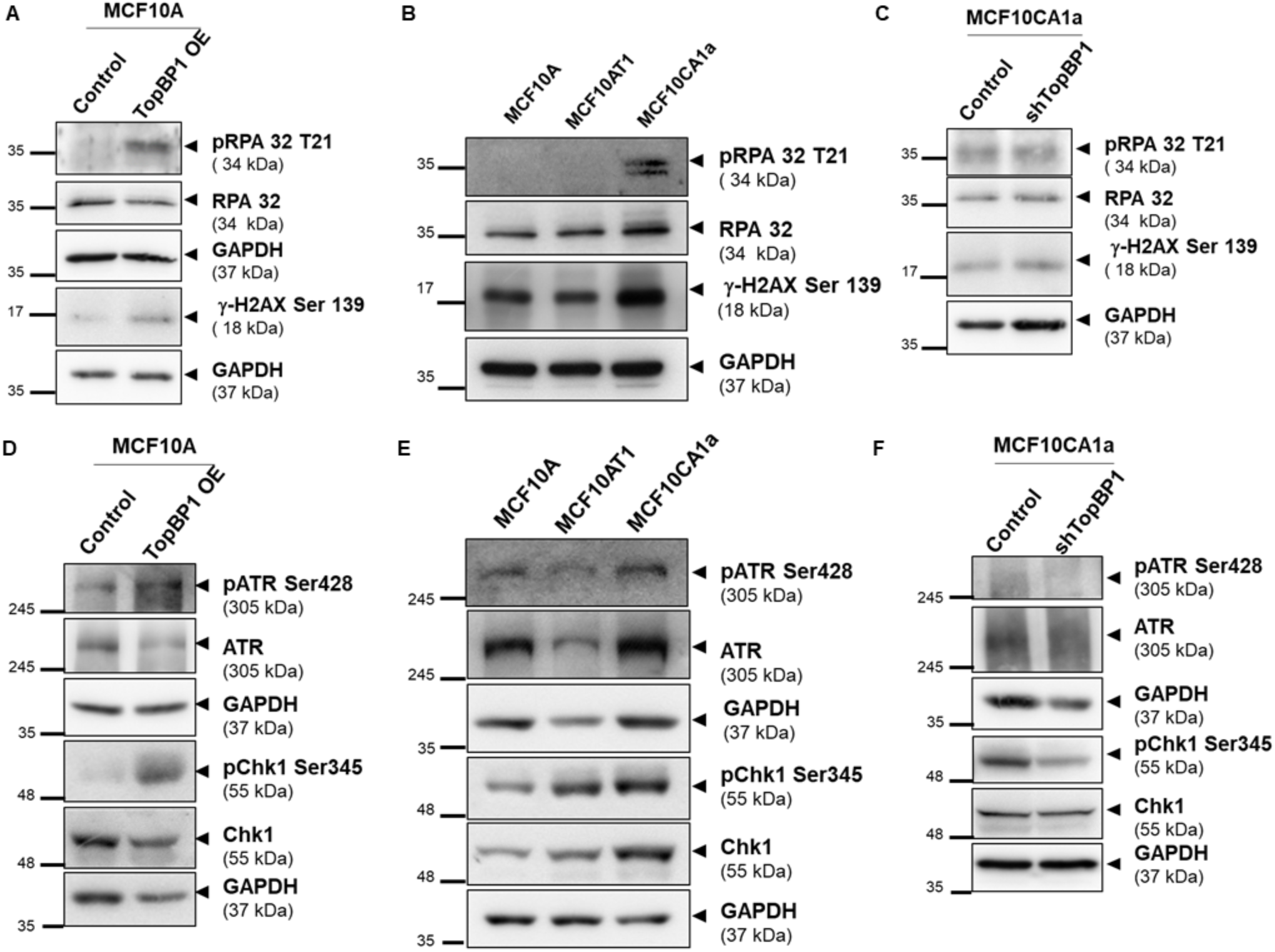
TopBP1 OE induces DNA damage and DDR activation. A and D) mCherry MCF10A (control) and TopBP1 OE MCF10A 3D dissociated cells, B and E) MCF10A, MCF10AT1, MCF10CA1a (2D) and C and F) control (pLKO1 MCF10CA1a) and shTopBP1 MCF10CA1a (2D) cells were lyzed and immunoblotting was performed to analyze the levels/activation of A-C) RPA, γH2AX D-F) ATR and Chk1. The phosphoproteins were quantified after normalizing to GAPDH.

### TopBP1 overexpression leads to TP53 mutation

p53 is a well-known tumor suppressor gene, which is an effector protein in the DNA damage checkpoint signaling pathway. Upon checkpoint activation, p53 is activated leading to a temporary cell cycle arrest to allow for DNA repair to occur. If the damage is beyond repair apoptosis signaling cascade is activated. In order to investigate whether TopBP1 overexpression leads to activation of p53 to execute a checkpoint response, total and activated p53 levels were analyzed in TopBP1 OE cells. Significant increases in total p53 and phosphorylated p53 at serine 15 were observed in TopBP1 OE cells when compared to the control cells (Figure 9A-C). Similar increases in total protein levels of p53 and phosphorylated p53 at serine 15 were also observed in MCF10CA1a cells in comparison to the pre-malignant MCF10AT1 and non-tumorigenic MCF10A cells (Figure 9E-G). However, p21, a p53 target protein that is active during G1-S arrest, was observed to be reduced or absent in TopBP1 OE (Figure 9A and D) and malignant cells (Figure 9B and H). Knockdown of TopBP1 in MCF10CA1a cells led to a significant decrease in the activation and total levels of p53 (Figure 9I-K) with a concomitant increase in p21 levels (Figure 9I and L). Studies have reported overexpression of TopBP1 to induce increased proliferation and reduced apoptosis through mutant p53 (mut p53) gain of function (Liu et al., 2011). Earlier reports have shown that polarity disruption in MCF10A cells can be induced by mutant p53 gain of function by inducing EMT (Zhang et al., 2011). Thus, TP53 mutation status was analyzed in mCherry MCF10A (control), TopBP1 OE MCF10A 3D dissociated cells and MCF10CA1a cells using whole-exome sequencing. Interestingly one mutation at 913 cDNA position in the TP53 gene was observed in TopBP1 OE cells, which was not present in the control cells (Table 1). This single-nucleotide variation (snv) mutation led to an exonic missense variant leading to the formation of a mutant p53 protein with isoleucine at 237 instead of methionine (M237I). This M237I mutation resides in the DNA binding region of p53 that is required for p53-TopBP1 interaction. This exact M237I mutation was also observed in the malignant cell line MCF10CA1a that has been demonstrated to have increased levels of TopBP1 suggesting that TopBP1 overexpression can also regulate the genomic stability inside the cells.

**Figure 9:**
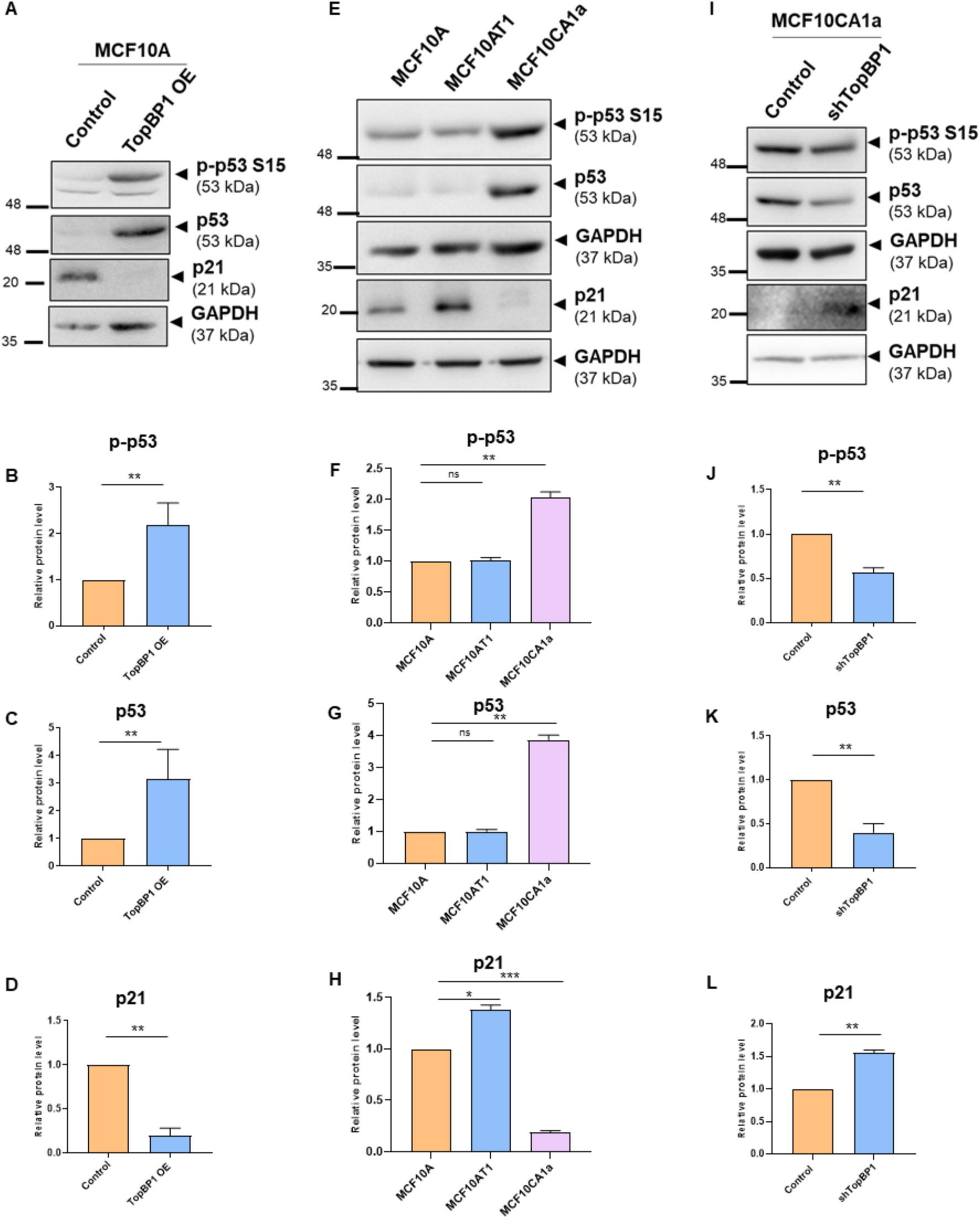
TopBP1 OE induces aberrant p53 activation. A) mCherry MCF10A (control) and TopBP1 OE MCF10A 3D dissociated cells, E) MCF10A, MCF10AT1, MCF10CA1a (2D) and I) pLKO1 MCF10CA1 (control) and shTopBP1 MCF10CA1a (2D) cells were lyzed and immunoblotting was performed to analyze the levels/activation of p53 and p21. Phosphoproteins were quantified after normalizing to GAPDH. Quantification showing the fold change of B, F, J) p-p53, C, G, K) p53 and D, H, L) p21 after normalizing to GAPDH. Statistical analysis was done using paired Paired Student’s t-test for TopBP1 OE and shTopBP1 lysates and One-way ANOVA for MCF10A series (*** p <0.001, ** p <0.01, * p <0.05, ns p>0.05).

**Table 1:**
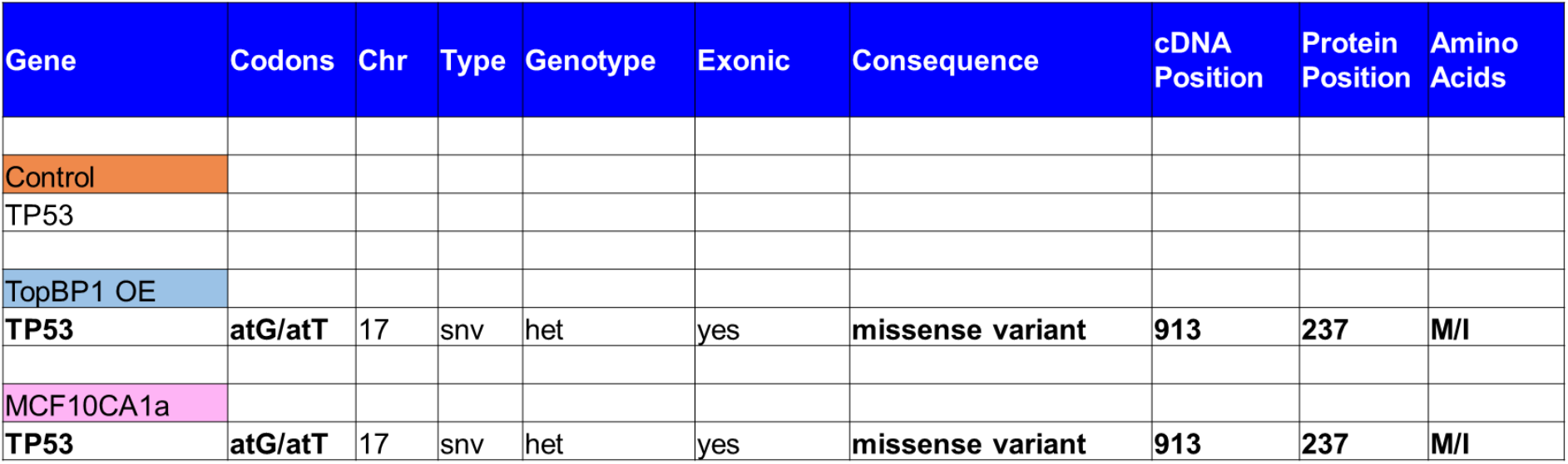
TP53 mutation in TopBP1 OE and MCF10CA1a cells

| Gene | Codons | Chr | Type | Genotype | Exonic | Consequence | cDNA Position | Protein Position | Amino Acids |
| --- | --- | --- | --- | --- | --- | --- | --- | --- | --- |
| Control |  |  |  |  |  |  |  |  |  |
| TP53 |  |  |  |  |  |  |  |  |  |
| TopBP1 OE |  |  |  |  |  |  |  |  |  |
| TP53 | atG/atT | 17 | snv | het | yes | missense variant | 913 | 237 | M/I |
| MCF10CA1a |  |  |  |  |  |  |  |  |  |
| TP53 | atG/atT | 17 | snv | het | yes | missense variant | 913 | 237 | M/I |

## Discussion

Cancer is caused by mutations in genes, which may give rise to increased cellular proliferation, decreased apoptosis, increased cell migration and invasion and/or EMT (Hanahan and Weinberg, 2011). These mutations in the genes are caused by various mutagens and carcinogens that work in favor of transforming the cells into a cancerous phenotype.

TopBP1 has been reported as a breast and ovarian cancer susceptibility gene by various groups (Forma et al., 2012; Karppinen et al., 2006). TopBP1 upregulation has been reported in high-grade breast cancer biopsy samples (Forma et al., 2012; Going et al., 2007; Liu et al., 2009).

In our study, we observed increased protein expression of TopBP1 in the malignant cells compared to the non-tumorigenic breast epithelial cells. We demonstrate that TopBP1 overexpression can induce transformation of non-tumorigenic MCF10A 3D breast acinar cultures to acquire properties similar to that of the malignant MCF10CA1a cells by inducing genomic instability and mutation in the TP53 gene (Figure 10). Our findings also emphasize the functional role of the cellular microenvironment in cancer initiation where TopBP1 overexpressing cells grown as 3-dimensional cultures underwent cellular transformation, which is not feasible by the cells when grown as monolayer cultures.

**Figure 10:**
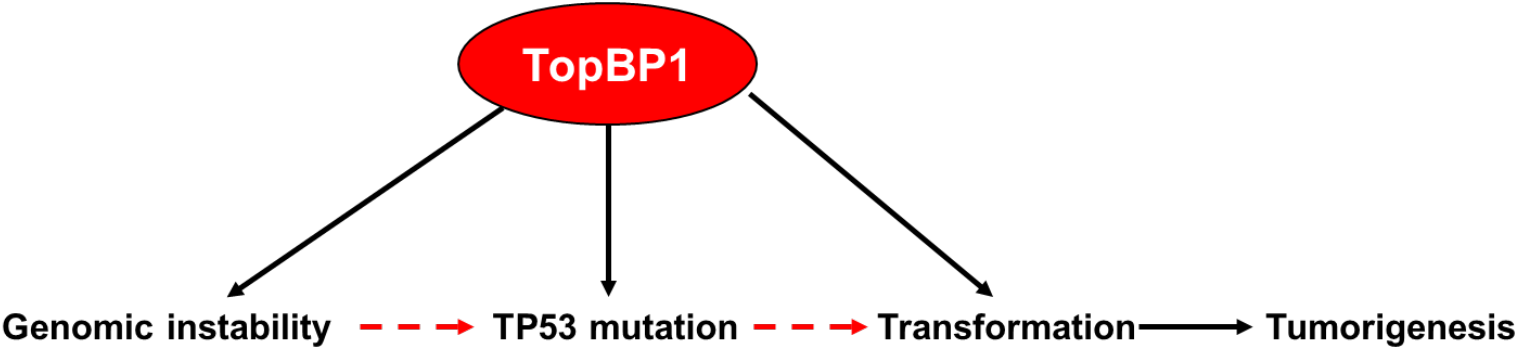
Schematic representation depicting the role of TopBP1 tumorigenesis. Overexpression of TopBP1 led to TP53 mutation in MCF10A cells grown as breast acinar cultures. This suggests that TopBP1 has the potential to regulate genome stability. This genome stability regulatory function of TopBP1 is also context dependent as TP53 mutation was not observed in TopBP1 overexpressing cells grown as monolayer cultures. This also emphasizes the role of the 3D environment in the transformation process as a change in morphology was not observed in MCF10A cells overexpressing TopBP1 grown as monolayer cultures.

Analysis of the cellular and nuclear pleomorphism and extent of the architectural disorder are the preliminary diagnostic and prognosis determination method for breast carcinoma (Debnath and Brugge, 2005). 3D cultures of breast epithelial cells and cancer cells recapitulate most of the features of these cells *in vivo*. Detailed analysis of these characteristic features can aid in ascertaining the role of perturbed genes. The morphometric analysis of TopBP1 OE acini revealed that TopBP1 overexpression induced changes in the acinar morphology which resembled cancerous spheroids. TopBP1 overexpressing cells formed spheroids with a larger surface area and volume and with filled lumen. In support of this observation, TopBP1 is known to play a major role in maintaining genome stability, ensuring proper DNA replication and G1-S cell cycle transition (Kim et al., 2005); perturbation in its level can distort DNA synthesis and cell cycle which may in turn, alter the nuclear content. We also found that TopBP1 OE acini to have increased proliferation, with a greater percentage of cells in the S and G2-M phases of the cell cycle in comparison to the MCF10A control acini.

De-regulation of apicobasal polarity is one of the characteristics of epithelial cancers (Debnath and Brugge, 2005). In invasive cancers, cells lose contact with neighbouring cells and break down the basement membrane to invade other tissues (Moreno-Bueno et al., 2008). In our study, we demonstrate TopBP1 overexpression leads to disruption of basal polarity. The loss of basal polarity markers a6-integrin and Laminin V indicated reduced interaction of the acinar cells with the ECM, which may indicate an invasive phenotype. We also found that overexpression of TopBP1 leads to Golgi dispersal, a known apical marker in MCF10A 3D cultures. Golgi dispersal is observed in various cancer cell lines, which may be the reason for altered polarity as Golgi plays a major role in maintaining polarized trafficking of proteins (Griffiths and Simons, 1986; Keller et al., 2001).

EMT is a process through which epithelial cells de-differentiate and acquire mesenchymal phenotype that can increase their motility and invasiveness. However, recent studies suggest that partial EMT phenotypes help in migration and metastasis (Aiello et al., 2018). We observed a similar pattern of expression of various EMT markers between TopBP1 OE and MCF10CA1a cells. N-cadherin and vimentin (mesenchymal markers) were downregulated, while E-cadherin (an epithelial marker) was upregulated in TopBP1 OE dissociated cells which was similar to that observed in MCF10CA1a cells. A recent study also suggests the role of E-cadherin in promoting metastasis in invasive ductal carcinoma (Padmanaban et al., 2019),which changes our perspective on EMT in tumorigenesis. This may hold true in our scenario as overexpression of TopBP1 led to increased migration and invasion. MCF10A cells are non-tumorigenic breast epithelial cells that require cell-cell contact and anchorage for their growth and do not have tumor forming ability. We observed that TopBP1 OE cells induced colony formation, anchorage-independent growth and tumorigenesis, thus confirming the role of TopBP1 as a potent tumor initiator and progression factor.

ATR is a major sensor kinase that gets activated by single-strand breaks and leads to activation of Chk1 to induce cell cycle arrest and DNA repair. TopBP1 is an important mediator protein in this signaling pathway. Even though TopBP1 levels have been observed to be high in cancer cells, its role in inducing DNA damage was not proven. A recent study from Lin’s group suggested an association between TopBP1 overexpression and genomic instability by TCGA analysis (Liu et al., 2021).We demonstrated that TopBP1 OE induced DNA damage and activation of DNA damage repair proteins in non-tumorigenic cells. We also observed that malignant cells have endogenous DNA damage compared to non-tumorigenic cells and altered activation of the proteins involved in the DNA damage response. Also, TopBP1 knockdown was able to partially revert these phenotypes thus confirming the role of TopBP1 in inducing genome instability. p53 is a well-studied tumor suppressor gene that is generally mutated in cancers. TopBP1 is known to bind to the DNA binding domain of p53 and inhibit its transcription activity of inducing apoptosis and cell cycle arrest without affecting its activation at S15 and total protein levels (Liu et al., 2009). However, we observed total p53 and p53 S15 levels were significantly increased in TopBP1 OE 3D dissociated cells. Even though p53 levels were high, p21 levels were low, as well as the proportion of cells in S and G2-M phases were high in TopBP1 OE dissociated cells confirming the inability of p53 to induce cell cycle arrest. Lin’s group have also demonstrated the role of TopBP1 in inducing mutant p53 gain of function to induce proliferation and reduce apoptosis (Liu et al., 2011). It was surprising that TopBP1 OE in a non-tumorigenic MCF10A cell line grown for 16 days in 3D culture had a similar TP53 mutation as was observed in the malignant cell line MCF10CA1a that was derived from MCF10A following H-Ras mutation and then passaging for 100 generations in mice. This TP53 mutation led to the formation of the mut p53 protein with an amino acid change from methionine to isoleucine at the 237^th^ reside (M237I). This mutation is reported in various cancers, where the somatic mutation has been reported in 176 cancers, out of which 26 were breast cancers (Jordan et al., 2010). Interestingly, SUM149PT, which is a triple negative breast cancer (TNBC) cell line harbors a p53 mutation at M237I (Wasielewski et al., 2006). M237I-p53 forms amyloid oligomers in glioblastoma cells expressing a chemoresistant gain-of-function phenotype (Pedrote et al., 2020).

Thus taken together, overexpression of TopBP1 induces transformation of non-tumorigenic breast epithelial cells when grown on extracellular matrix as 3-dimensional cultures. These transformed cells have phenotypic and genotypic characteristics concurrent to malignant cells with a mutation in the TP53 gene. Our study also opens multiple avenues for investigating and tracking the initiation as well as early-stage changes in cancers. TopBP1 being a novel regulator in the process of tumorigenesis, also exposes itself as a potential target to combat cancer(s). Cancers resulting from higher TopBP1 protein expression can be targeted with therapeutic molecules that can either inhibit the function or reduce the half-life of TopBP1 inside cells, thus eventually regulating genomic instability and functions of p53.

## Materials and Methods

### Ethics statements

Institutional Biosafety Committee (IBSC) clearance was obtained from the institute for preparing TopBP1 OE MCF10A cells and TopBP1 KD MCF10CA1a cells. All mouse studies were approved by the Institutional Animal Ethics Committee (IAEC) (IAEC/2018_02/010 and IISER_Pune/IAEC/2021_01/06). We only used athymic mice (Foxnl ^nu^ / Foxnl ^nu^, 6-8 weeks old) for the *in vivo* tumorigenicity studies

### Chemicals and antibodies

Insulin (I1882), Hydrocortisone (H0888), Choleratoxin (C8052), Epidermal growth factor (EGF-E9644), Puromycine dihydrochloride from Streptomyces alboniger (P8833), Polybrene (H9268), dimethyl sulfoxide (DMSO, D8418), Triton X-100 (T8787), Mitomycin C (M4287), Tris base (B9754), EDTA (E6758), propidium iodide (PI-P4170) and Na3VO (S6508) were purchased from Sigma-Aldrich, USA. Matrigel®and Dispase (354235) were purchased from Corning, Sigma-Aldrich, USA. DQ Collagen (D12060), Phalloidin 633 (A12380) and Lipofectamine-2000 (11668) were purchased from Invitrogen, Thermo Fisher Scientific, USA. Pre-stained protein ladder (MBT092), NaF (RM1081), Na_2_HPO_4_ (GRM1417), and KH_2_PO_4_ (MB050) were purchased from HiMedia. NaCl (15918), was purchased from Qualigens. 16% paraformaldehyde (AA433689M) was purchased from Alfa Aesar. Hoechst 33258 (H3569) and Hoechst 33342 (H3570) were purchased from Molecular Probes, Thermo Fisher Scientific, USA. Antibodies used were from Bethyl, Abcam, Millipore, cell signalling (CST), Santacruz or Sigma (Supplementary Table 1).

### Cell lines

HEK293T and MCF10A were generous gifts from Dr Jomon Joseph (NCCS, Pune) and Prof. Raymond C. Stevens (The Scripps Research Institute, California) respectively. MCF10AT1 and MCF10CA1a cells were purchased from ATCC.

### Plasmids

CSII-EF-MCS plasmid was a gift from Dr Sourav Banerjee, NBRC, Manesar India. pCAG-HIVgp and pCMV-VSV-G-RSV-Rev plasmids were purchased from RIKEN BioResource Centre, Japan. pLKO.1.TRC was a gift from Dr Sorab Dalal, ACTREC, India and psPAX2 and pMD2.G were generous gifts from Dr Manas K Santra, NCCS, India. The vector maps are provided in Supplementary Figures S3-S6.

### Cloning

TopBP1 was amplified from mVenus TopBP1 vector using the following primers: Forward: AAGGAAAAAAGCGGCCGCATATGTCCAGAAATGACAAAGAACCG Reverse: ATGGTTAACTTAGTGTACTCTAGGTCGTTTGATT (PCR mixture and reaction in Supplementary Table 2A-B). The amplified plasmids and CSII-EF MCS-mCherry vector were digested using HpaI and NotI HF (NEB, USA) restriction enzymes (Supplementary Table 2C). TopBP1 was ligated into mCherry-CSII-EF-MCS vector using T4 DNA ligase (NEB, USA) following incubation at 16°C overnight (Supplementary Table 2D). Ligation mixture was used to transform DH5α competent cells, and the colonies were screened by digesting with HpaI and NotI HF to check for insert release.

shTopBP1 lentiviral plasmids were generated as per Addgene protocol. Briefly, pLKO1. TRC vector was digested with EcoRI and AgeI restriction enzymes (NEB, USA) (Supplementary Table 3A), and the 7 kb fragment was excised and gel extracted using QIAquick Gel Extraction Kit (28706). The shTopBP1 oligos (forwad: CCGGAAGTGGTTGTAACAGCGCATCTTCTCGAGAAGATGCGCTGTTACAACCACTTTT TTTG and reverse: AATTCAAAAAAAGTGGTTGTAACAGCGCATCTTCTCGAGAAGATGCGCTGTTACAACC ACTT) were annealed (Supplementary Tables 3B-C), and ligated into the digested vector (Supplementary Table 3D). The clones obtained were screened using XhoI (NEB, USA) digestion.

### Cell culture

HEK293T cells were grown on 100mm tissue culture-treated dishes (Eppendorf or Corning) with high glucose-containing Dulbecco’s Modified Eagle Medium (DMEM; Lonza) supplemented with 10% heat-inactivated fetal bovine serum (Invitrogen, Thermo Fisher Scientific, USA) and 100 units/ml of penicillin-streptomycin (Invitrogen, Thermo Fisher Scientific, USA). MCF10A, MCF10AT1 and MCF10CA1a cells were grown as monolayer cultures in growth medium and 3D in assay medium as previously mentioned (Anandi et al, 2017). Cells were incubated at 37°C in humidified 5% CO2 incubators (Eppendorf or Thermo Fisher Scientific, USA).

### Lentiviral preparation and transduction using lipofectamine-mediated transfection

Lentiviral particles to generate TopBP1 overexpressing MCF10A cells were prepared as previously mentioned (Sharma and Lahiri, 2021). Briefly, HEK293T cells were transfected with 1 μg of mCherry-CSII-EF-MCS or mCherry-CSII-EF-MCS-TopBP1 plasmid along with 1 μg of pCAG-HIVgp and 0.5 μg pCMV-VSV-G-RSV-Rev packaging plasmids using Lipofectamine 2000. 1ml growth media containing 15% horse serum was added to the cells 24 hours post-transfection. The viral supernatant collected 48 hours post-transfection was used to transduce MCF10A cells. Transduced MCF10A cells were sorted using the BD Aria Fusion sorter (BD Biosciences, USA).

For the generation of MCF10CA1a cells with TopBP1 knockdown, HEK293T cells were transfected with 1 μg of pLKO1.TRC or shTopBP1-TRC plasmid along with 1 μg of psPAX2 and 0.5 μg pMD2 packaging plasmids using Lipofectamine 2000 to generate lentiviral particles. These lentiviral particles were used to transduce MCF10CA1a cells, and cells with TopBP1 knockdown were selected using 1 μg/ml puromycin.

### 3D dissociation

For dissociating cells from 3D acini, the 16-day 3D cultures were treated with 250 μl of Dispase and incubated at 37°C for 30 minutes. The dissociated cells along with Matrigel®were transferred to a 15 ml falcon and incubated at 37°C for 10 minutes. The dislodged acini were centrifuged at 900 rpm for 10 minutes and plated in 12-well culture plates after washing twice with 1X PBS. The attached cells were passaged and expanded once 70% confluency was reached.

### Immunofluorescence

Immunofluorescence of 2D and 3D cultures were performed as mentioned previously (Anandi et al., 2017). For IF of 3D cultures, 0.5 % Triton X was added to the wash buffers for better penetration of antibodies. Cells / acini were imaged on the SP8 confocal microscope (Leica, Germany) under 63X/40X oil immersion objective. A minimum of 200 cells or 50 acini were analyzed for the phenotypic characterization.

### Immunoblotting

Immunoblotting analysis using specific antibodies were performed as mentioned previously (Bodakuntla et al., 2014). Images were acquired on ImageQuant LAS4000 gel documentation system (GE Healthcare, now Cytiva Life Sciences).

### Cell proliferation assay

For cell proliferation assay, 2.5 × 10^5^ mCherry or TopBP1 OE MCF10A 3D dissociated cells were seeded on a 35mm dish. 72 hrs post seeding, the cell count was measured using Sceptar™ hand-held automated cell counter (Millipore, Sigma-Aldrich, USA) as previously mentioned (Sharma and Lahiri, 2021).

### Flow cytometry

The cell cycle profile was analyzed as mentioned previously (Sharma and Lahiri, 2021). Cell cycle profile was analyzed on a BD Accuri™ C6 flow cytometer (BD Biosciences, USA).

### Wound healing Assay

Wound healing analysis was performed as mentioned previously (Anandi et al., 2016). Images were taken at 0 and 12 hrs of the assay at 10X objective on the ECLIPSE TS100 (Nikon, Japan) microscope and wound area was manually measured using ImageJ software (NIH, USA)

### Single-cell migration assay

Cells were seeded in 24-well cell culture plates at a density of 2 × 10^5^ cells per well and incubated at 37°C for 10 hrs. Cells were stained with Hoechst 33342 for 15 minutes and replenished with fresh L15 media (12-700Q, Lonza). Cells were tracked for 3 hrs at 37°C, and images were taken at every 2 mins time interval on the EVOS FL Auto microscope (Life Technologies) at 20X magnification. The cells were tracked using Fast track software (DuChez, 2017), and the distance, displacement and velocity obtained were plotted in Graphpad prism (Graph Pad Software, La Jolla, CA, USA).

### Gelatin Zymography

Assay medium of 3D cultures were collected 10 days post seeding and mixed with gelatin zymography sample buffer (25mM Tris-HCl, pH 7.6, 1mM NaCl, 1% NP-40). It was then resolved on a 8% SDS-PAGE containing 0.06% gelatin [w/v] as substrate. The gel was renatured with 1X renaturing buffer (2.5 % v/v of Triton X-100 in water) at room temperature (RT) for 30 minutes with constant shaking. After washing with distilled water, the gel was incubated in 1X Developing buffer (0.5 M Tris-HCL pH-7.8, 2M NaCl, 0.05M CaCl2 and 0.2% Brij35) for 48 hours. Meanwhile the same media mixed with 6X sample buffer was resolved on a 8% SDS-PAGE to find the total protein content in the media. Later, the gels were stained overnight in staining solution (0.1% w/v Coomassie Brilliant Blue (Sigma-Aldrich), 50% v/v methanol (Thermo Fisher Scientific, USA) and 10% v/v acetic acid (Thermo Fisher Scientific, USA) and destained with destaining solution (40% v/v methanol and 10% v/v acetic acid) for 1 hour. The gel images were acquired on ImageQuant LAS4000 gel documentation system (GE Healthcare now Cytiva Life Sciences).

### Colony formation assay

Cells were seeded in 6-well culture plates at a density of 300 cells per well. Cells were replenished with fresh media after 4 days of seeding. After 7 days of seeding, cells were washed twice with 1X PBS and fixed with 100% chilled methanol at 4°C for 15 minutes. Cells were stained with 0.5% of crystal violet stain (Himedia, S012) for 15 minutes with shaking at RT. After removing the stain, cells were washed with distilled water and images were taken on a HP Scanner (HP DeskJet 2332) after drying the plates. The number of colonies were manually counted.

### Soft agar assay

Soft agar assay was performed as mentioned previously (Anandi et al., 2017). Ten randomly selected fields were imaged under 10X objective of ECLIPSE TS100 (Nikon, Japan) microscope and number of colonies were counted manually.

### Tumorigenesis in mice

For mice injections, 6 × 10^6^ mCherry or TopBP1 MCF10A 3D dissociated cells were resuspended in 100 μl of 1:1 mixture of PBS: Matrigel and injected into the flanks of athymic nude mice (Foxnl ^nu^ / Foxnl ^nu^, 6-8 weeks old) (IAEC/2018_02/010). The mice were examined every week. The mice were sacrificed on the 7^th^ week and the tumors were dissected. The tumor size and volume were calculated by measuring the tumor dimensions using a Vernier calliper.

### Whole exome sequencing

Genomic DNA was extracted from mCherry-MCF10A, TopBP1 OE 3D dissociated and MCF10CA1a cells using the DNeasy blood and Tissue kit (Qiagen, 69506) as per manufactures protocol. All three cell lines were sent to Dhiti Omics Technologies, Bengaluru, India for whole exome sequencing (Makiniemi et al.) using Agilent V6r2 (Agilent Technologies, Santa Clara, CA 95051, USA) and this targeted library was sequenced using HiSeq. The resultant raw data were then mapped to the human genome (Hg19/GRCh37) using BWA algorithm (Abuin et al., 2015; Li and Durbin, 2009, 2010) and duplicates were removed using Picard toolkit (Github, Board institute). For all 3 cell lines mapping were in the cut-off limit (i.e mapping= >95% and Read Duplicates <20%). Further target enrichment was done using samtools and subsequent variant calling was done on BAM file using GATK pipeline to produce VCF formatted output file (BSD, MIT). All variants in the VCF file were then annotated using in-house pipeline with all required databases including OMIM, ClinVar HGBS coordinates, different population databases, etc (Ulintz et al., 2019). Sequencing data was analyzed manually to identify TP53 gene mutations in the cell lines.

### Statistical Analysis

At least 3 biological independent replicates were performed for all the experiments. Hygens software (SVI, Hilversum, Netherlands) was used to calculate the surface area, volume and sphericity of the 3D acini or spheroids. Results were analyzed and plotted using GraphPad Prism (GraphPad Software, La Jolla, CA, USA). Mann Whitney U test was used to test the significance of difference of various parameters in the morphometric analysis and immunofluorescence. Paired Student’s t-test or One-way ANOVA was performed to analyze the significance of difference of protein expression in immunoblotting.

## Author Contributions

R.M.U and M.L conceived and conceptualized the project. R.M.U designed, performed and analyzed the experiments. R.M.U and M.L wrote the paper.

## Acknowledgements

We thank Drs Richa Rikhy (IISER, Pune, India), and Sorab Dalal (ACTREC, Navi Mumbai, India) for their valuable suggestions. The authors acknowledge the IISER Pune Microscopy Facility and IISER-BD FACS facility for access to equipments and infrastructure. We thank National Facility for Gene Function in Health and Disease (IISER, Pune, India) for providing the animals and facilities. We also thank Lahiri lab members for their helpful comments and discussions.

