## Supplementary Information for "Overexpression of TopBP1 leads to transformation with a TP53 mutation of non-tumorigenic breast epithelial cells"

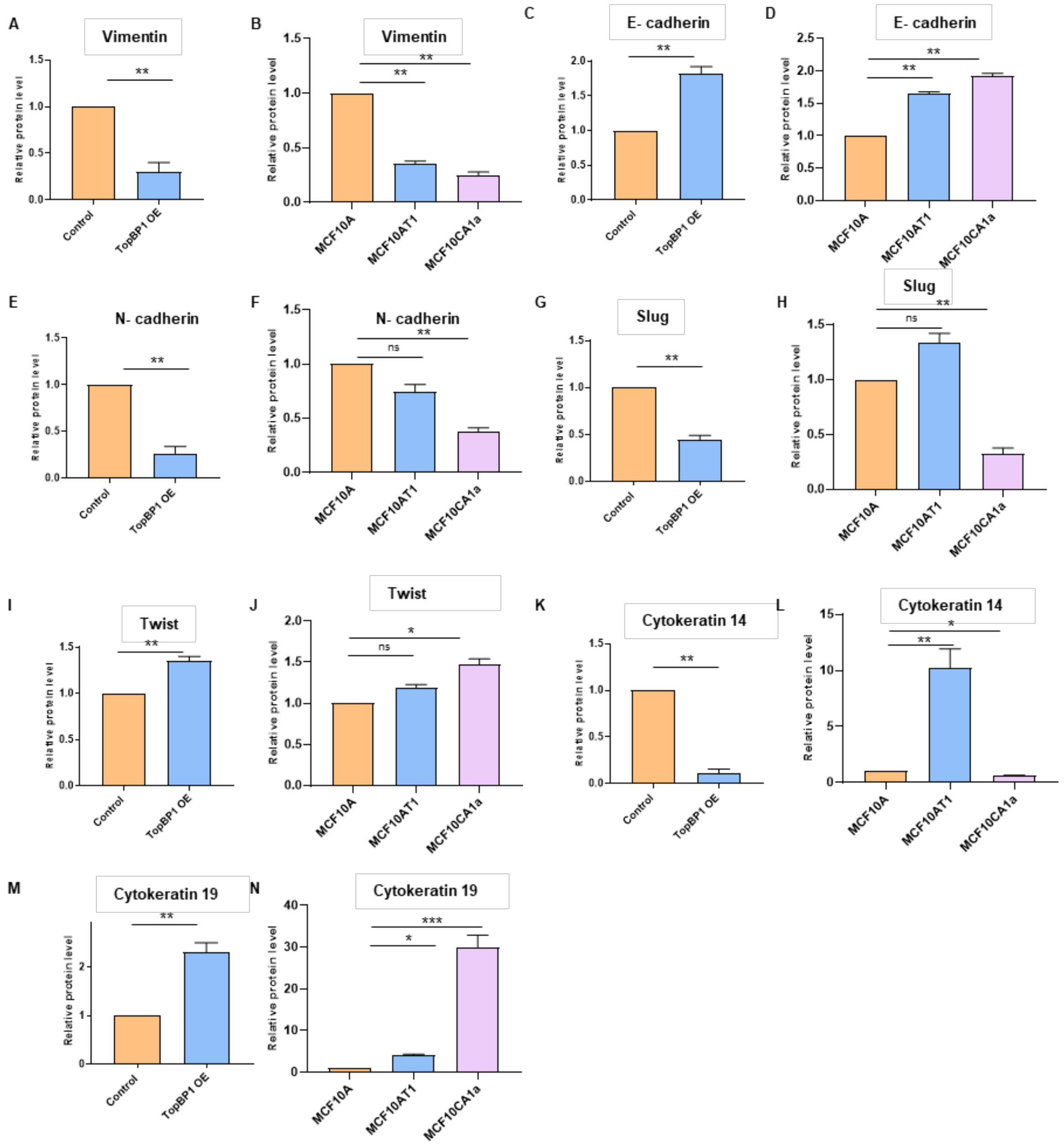


**Supplementary Figure S1:** To analyze the protein expression of various EMT markers, the cells were lyzed and immunoblotting was performed. A, C, E, G, I, K, M) mCherry MCF10A (control) and TopBP1 OE MCF10A 3D dissociated cells and B, D, F, H, J, L, N) MCF10A, MCF10AT1 and MCF10CA1a cells. Quantification was done to show the fold change of A-B) vimentin, C-D) E-cadherin, E-F) N-cadherin, G-H) Slug, I-J) Twist, K-L) Cytokeratin 14 and M-N) Cytokeratin 19 between mCherry MCF10A (control) and TopBP1 OE MCF10A 3D dissociated cells and MCF10A, MCF10AT1 and MCF10CA1a cell lysates after normalizing with the GAPDH. Statistical analysis was done using Paired Student's t-test for TopBP1 OE lysates and One-way ANOVA for MCF10A series (*** p <0.001, ** p <0.01, * p <0.05, ns p>0.05).

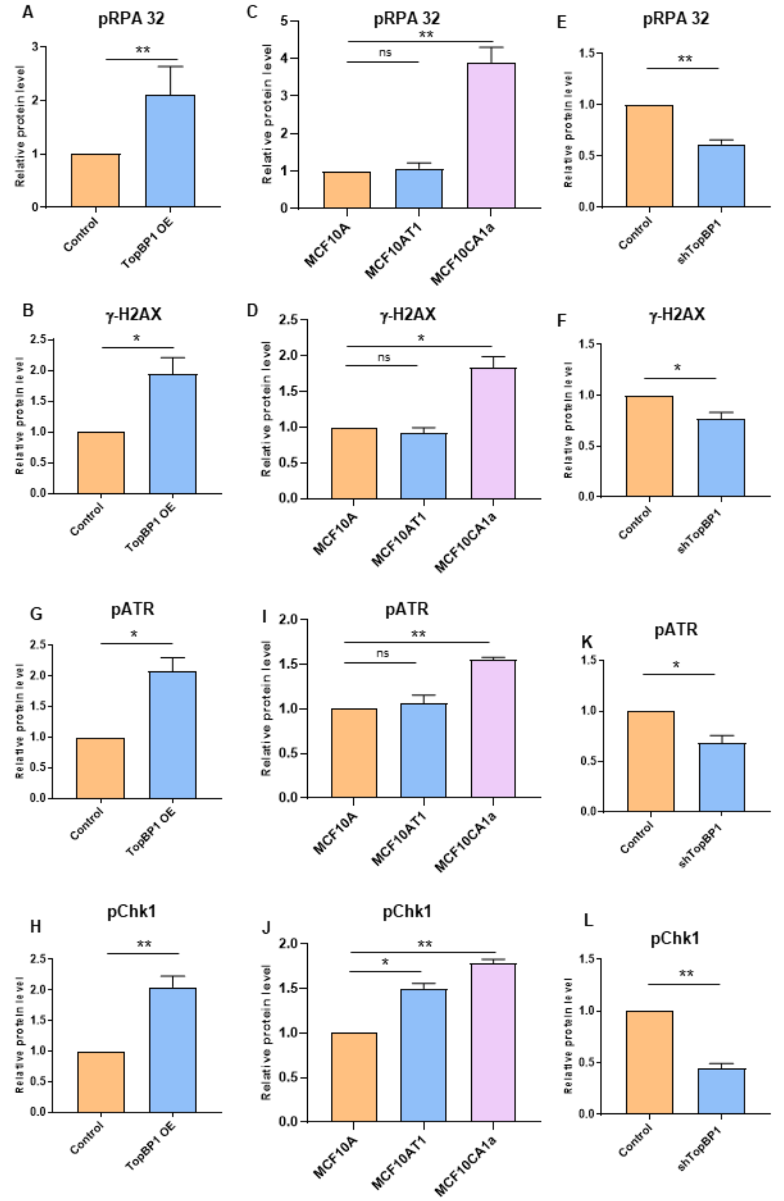


**Supplementary Figure S2:** To analyze the activation of various DNA damage repair proteins, the cells were lyzed and immunoblotting was performed A, B, G, H) mCherry MCF10A (control) and TopBP1 OE MCF10A 3D dissociated cells, C, D, I J) MCF10A, MCF10AT1, MCF10CA1a cells and pLKO.1 MCF10CA1a (control) and E, F, K, L) shTopBP1 MCF10CA1a cells. Quantification showing the fold change of A, C, E) pRPA 32, B, D, F) $\gamma$H2AX, G, I, K) pATR, and H, J, L) pChk1. Phosphoproteins were quantified after normalizing to GAPDH. Statistical analysis was done using Paired Student's t-test for TopBP1 OE and shTopBP1 lysates and One-way ANOVA for MCF10A series ( ** p <0.01, * p <0.05, ns p>0.05).

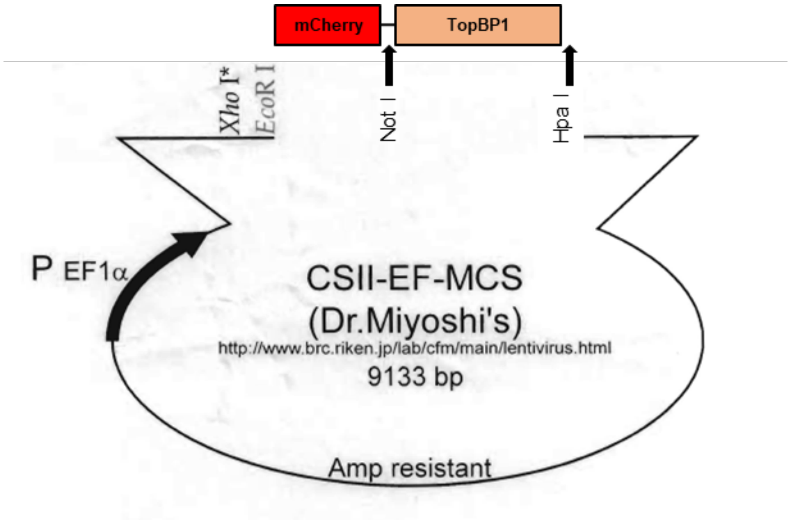


**Supplementary Figure S3:** TopBP1 was cloned into a CSII-EF-MCS vector with a mCherry tag between NotI and HpaI sites. The vector map of the CSII-EF-MCS plasmid was provided by Riken BioResource Centre, Japan.

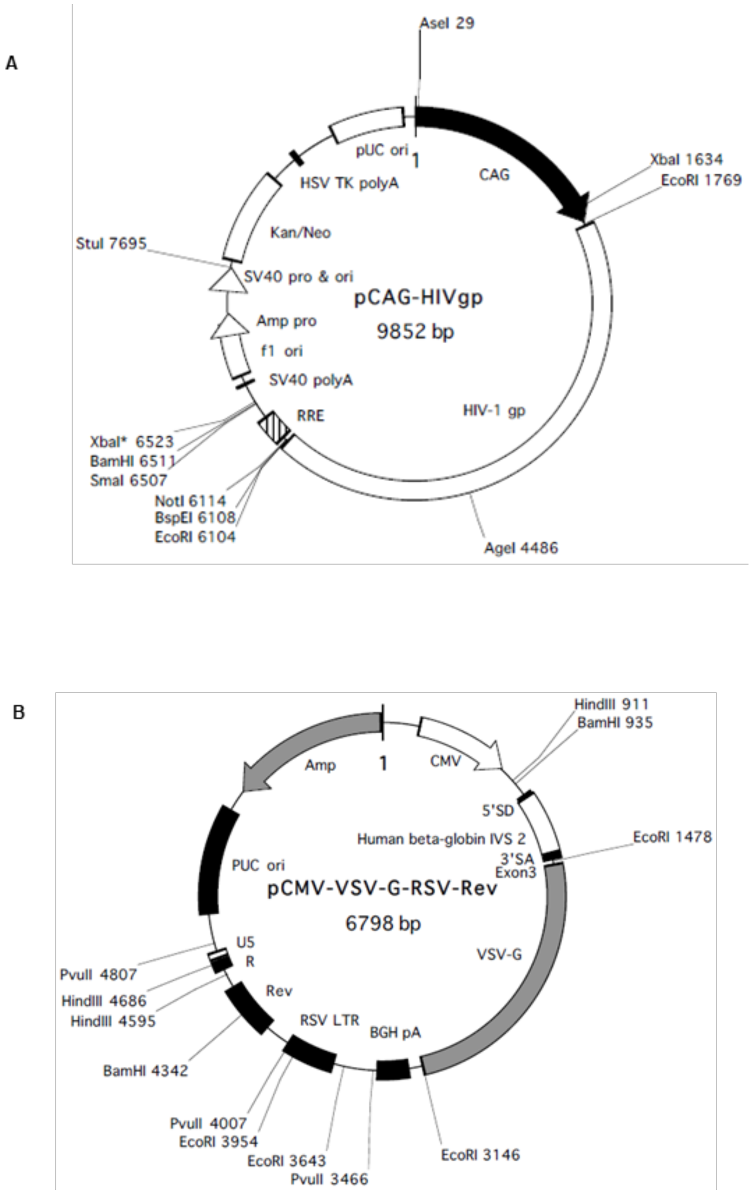


**Supplementary Figure S4:** Vector map of A) HIV-1 gag and pol for lentivirus vector packaging and B) envelope VSVG protein and Rev packaging plasmid for the generation of TopBP1 overexpression lentiviral particles. The plasmids was purchased from RIKEN BioResource Centre, Japan and the vector map was provided by them.

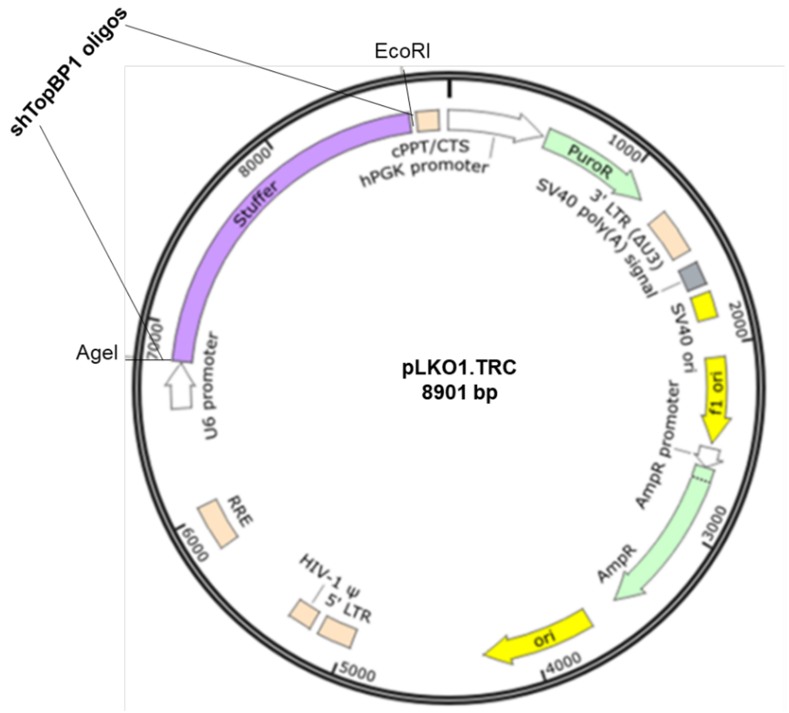


**Supplementary Figure S5:** shTopBP1 oligos were annealed and cloned into pLKO.1.TRC vector with a puro selection marker between AgeI and EcoRI sites. The plasmid was a generous gift from Dr Sorab Dalal, ACTREC, India. The vector map was obtained from Addgene web site

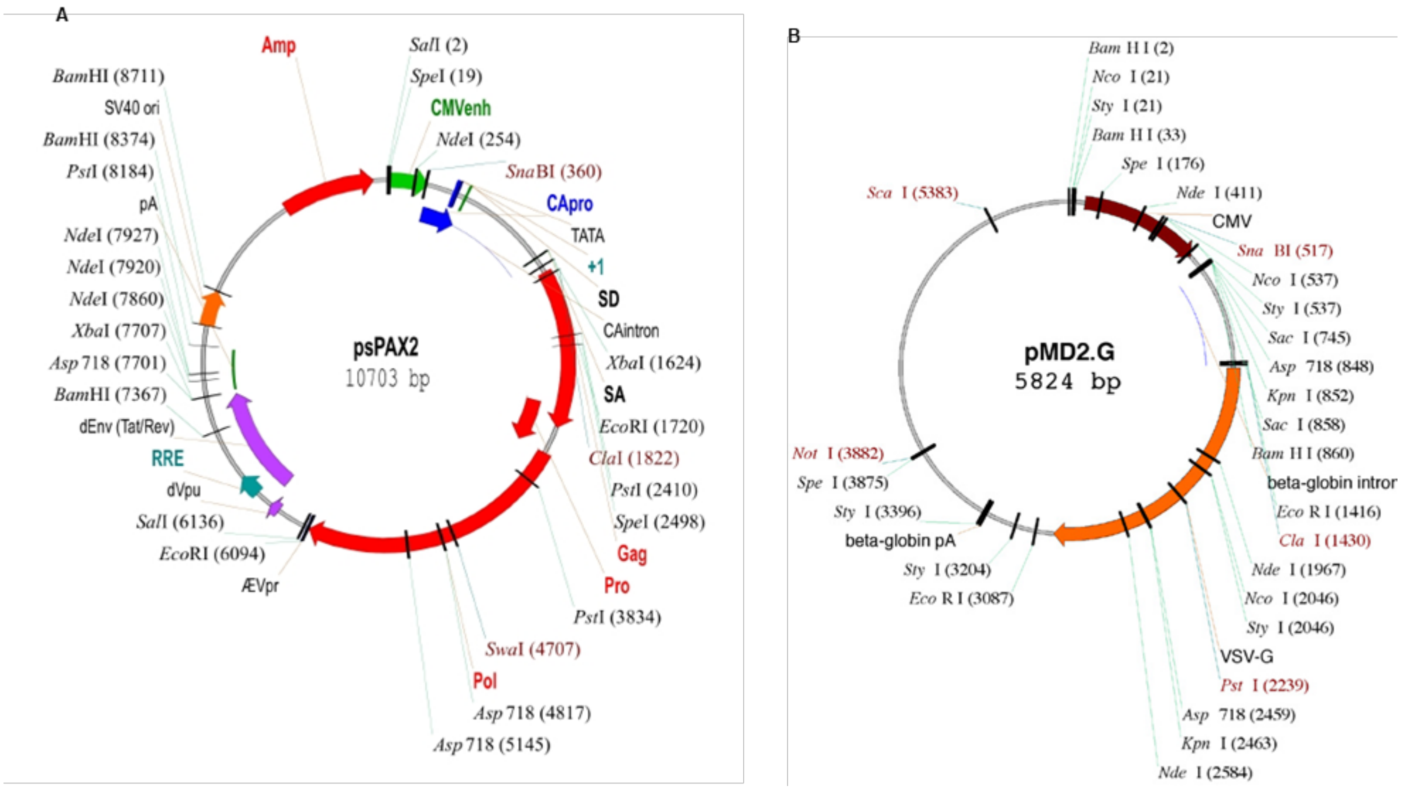


**Supplementary Figure S6:** Vector map of A) psPAX2 gag and pol for lentivirus vector packaging and B) epMD2.G VSVG protein packaging plasmid for the generation of shTopBP1 lentiviral particles. The plasmids were a generous gift from Dr Manas K Santra, NCCS, India. The vector maps were obtained from the Addgene website.

**Supplementary Table 1: Antibodies used for IF and Immunoblotting**

| *Primary Antibody* | Catalogue no | IF | Immunoblotting |
|---|---|---|---|
| RPA 32/2 Phospho T21 (Abcam) | AB 61065 | | 1:1000 |
| RPA (CST) | 2208 | | 1:1000 |
| Laminin V, anti-Mouse (Millipore) | MAB19562 | | |
| TopBP1, anti-Rabbit (Bethyl) | A300-111A | 1:200 | 1:1000 |
| Rat X Hu CD49fα6 integrin, anti –Rat (Millipore) | MAB1378 | 1:100 | - |
| GM130, anti-Mouse (Millipore) | 610822 | 1:100 | |
| E-cadherin, anti-Mouse (BD) | 610182 | | 1:7500 |
| N-cadherin, anti-Rabbit (Abcam) | AB18203 | | 1:1000 |
| Vimentin, anti-Mouse (Abcam) | AB 8069 | 1:200 | 1:10000 |
| β-catenin, anti- Rabbit (Abcam) | ab32572 | 1:100 | |
| PCNA- PC10 (CST) | 2586 | | 1:2500 |
| Cytokeratin 14, anti-Mouse (Abcam) | ab7800 | | 1:1000 |
| Cytokeratin 19, anti-Rabbit (Abcam) | ab52625 | | 1:1000 |
| GAPDH, anti-Rabbit (Sigma) | G9545 | | 1:40000 |
| Slug- C19G7 (CST) | 9585 | | 1:1000 |
| Twist (Abcam) | ab50581 | | 1:1000 |
| γH2AX Ser 139 (Santacruz) | sc517348 | | 1:1000 |
| pATR Ser 428 (CST) | 2853 | | 1:1000 |
| ATR (Abcam) | AB 2905-100 | | 1:1000 |
| pChk1 (CST) | 2348L | | 1:1000 |
| Chk1(Santacruz) | SC 8408 | | 1:1000 |
| p-p53 S15 (CST) | 9284S | | 1:1000 |
| p53 (Bethyl) | A300-247A | | 1:5000 |
| p21 (Abcam) | ab109520 | | 1:1000 |
| Cyclin A (Santacruz) | sc-271682 | | 1:1000 |

**Supplementary Table 2: Cloning of TopBP1 into CSII-EF MCS vector**

**A.** Amplification of TopBP1

| Reagents | Amount |
|---|---|
| DNA Template | 500 ng |
| Forward Primer | 3 μl |
| Reverse Primer | 3 μl |
| 10 mM dNTPs | 12.5 μl |
| 10X Pfu buffer | 10 μl |
| Pfu polymerase | 2 μl |
| Nuclease free water | To make up the volume to 100 μl |

**B.** PCR cycle for amplification of TopBP1

| Temperature (ºC) | Time (mins) | |
|---|---|---|
| 95 | 2 | |
| 95 | 1 | 25 cycles |
| 50 | 1 | 25 cycles |
| 72 | 10 | 25 cycles |
| 72 | 10 | |

**C.** Digestion of PCR product and CSII-EF-MCS-mCherry

| Digestion reagents | Amount |
|---|---|
| DNA Template | 5 μg |
| 10 X Cut smart buffer | 5 μl |
| Not I HF | 5 μl |
| Hpa I | 5 μl |
| Nuclease free water | To make up the reaction to 50 μl |

**D.** Ligation reaction of TopBP1 and CS-II-EF-MCS-mCherry digested vector

| Ligation reagents | Amount |
|---|---|
| Insert | 75 ng |
| Vector | 25 ng |
| 10X T4 Ligase buffer | 1 μl |
| T4 DNA Ligase | 1 μl |
| Nuclease free water | To make up the reaction to 10 μl |

**Supplementary Table 3: Cloning of shTopBP1 into pLKO1.TRC vector**

**A.** Restriction digestion of pLKO1.TRC vector

| Digestion Reagents | Amount |
|---|---|
| DNA | ~3 µg |
| 10x cut smart buffer (NEB) | 5 µl |
| AgeI Hf (NEB) | 3 µl |
| EcoRI Hf (NEB) | 3 µl |
| Nuclease free water | Make upto 50 µl |

**B.** Annealing reaction for shTopBP1 oligos

| Anealing Reagents | Volume (µl) |
|---|---|
| TopBP1 shRNA forward oligo | 1.0 |
| TopBP1 shRNA reverse oligo | 1.0 |
| 10 x T4 DNA Ligase buffer (Takara) | 1.0 |
| T4 PNK | 0.5 |
| Nuclease free water | 6.5 |

**C.** PCR program for annealing of shTopBP1 oligos

| Temperature | 37 °C | 95 °C | 90 °C | 85 °C | 80 °C | 75 °C | 70 °C | 65 °C | 60 °C | 55 °C | 50 °C | 45 °C | 40 °C | 35 °C | 30 °C | 25 °C | 4°C |
|---|---|---|---|---|---|---|---|---|---|---|---|---|---|---|---|---|---|
| Time | 30’ | 5’ | 1’ | 1’ | 1’ | 1’ | 1’ | 1’ | 1’ | 1’ | 1’ | 1’ | 1’ | 1’ | 1’ | 1’ | hold |

**D.** Ligation reaction for annealed shTopBP1 oligos and digested pLKO1.TRC

| Reagents | Amount |
|---|---|
| Digested pLKO.1-TRC | 25ng |
| shTopBP1 | 125ng |
| T4 DNA Ligase (Takara) | 1 µl |
| 10x T4DNA Ligase buffer(Takara) | 1 µl |
| Nuclease free water | Upto 10 µl |
